# Sleep loss drives brain region- and cell type-specific alterations in ribosome-associated transcripts involved in synaptic plasticity and cellular timekeeping

**DOI:** 10.1101/2020.07.20.212019

**Authors:** Carlos Puentes-Mestril, James Delorme, Marcus Donnelly, Donald Popke, Sha Jiang, Sara J. Aton

## Abstract

Sleep and sleep loss are thought to impact synaptic plasticity, and recent studies have shown that sleep and sleep deprivation (SD) differentially affect gene transcription and protein translation in the mammalian forebrain. However, much less is known regarding how sleep and SD affect these processes in different microcircuit elements within the hippocampus and neocortex - for example, in inhibitory vs. excitatory neurons. Here we use translating ribosome affinity purification (TRAP) and *in situ* hybridization to characterize the effects of sleep vs. SD on abundance of ribosome-associated transcripts in Camk2a-expressing (Camk2a+) pyramidal neurons and parvalbumin-expressing (PV+) interneurons in mouse hippocampus and neocortex. We find that while both Camk2a+ neurons and PV+ interneurons in neocortex show concurrent SD-driven increases in ribosome-associated transcripts for activity-regulated effectors of plasticity and transcriptional regulation, these transcripts are minimally affected by SD in hippocampus. Similarly we find that while SD alters several ribosome-associated transcripts involved in cellular timekeeping in neocortical Camk2a+ and PV+ neurons, effects on circadian clock transcripts in hippocampus are minimal, and restricted to Camk2a+ neurons. Taken together, our results indicate that SD effects on transcripts destined for translation are both cell type- and brain region-specific, and that these effects are substantially more pronounced in the neocortex than the hippocampus. We conclude that SD-driven alterations in the strength of synapses, excitatory-inhibitory balance, and cellular timekeeping are likely more heterogeneous than previously appreciated.

**Significance Statement:** Sleep loss-driven changes in transcript and protein abundance have been used as a means to better understand the function of sleep for the brain. Here we use translating ribosome affinity purification (TRAP) to characterize changes in abundance of ribosome-associated transcripts in excitatory and inhibitory neurons in mouse hippocampus and neocortex after a brief period of sleep or sleep loss. We show that these changes are not uniform, but are generally more pronounced in excitatory neurons than inhibitory neurons, and more pronounced in neocortex than in hippocampus.

## Introduction

Sleep is essential for optimal brain function, but the underlying biological mechanisms are largely unknown. Prior work aimed at addressing this question has used molecular profiling of mRNA and protein abundance, in a number of brain areas, to characterize changes caused by experimental SD (Cirelli et al., 2004; Mackiewicz et al., 2007; Noya et al., 2019; Poirrier et al., 2008; Vecsey et al., 2012). Transcriptomic changes reported after SD in the brain have led to specific hypotheses regarding the biological underpinnings of cognitive disruptions associated with sleep loss (Belenky et al., 2003; Dinges et al., 1997; Mednick et al., 2003; Stickgold, 2005). For example, the synaptic homeostasis hypothesis (Tononi and Cirelli, 2006) proposes that synapses throughout the brain are strengthened during periods of wake and weakened during periods of sleep. The proposal of this hypothesis was initially based on results from transcriptomic studies in mice, showing higher expression of both immediate early genes (IEGs) and several other genes involved in synaptic plasticity after periods of SD vs. sleep (Cirelli et al., 2004; Cirelli et al., 1996; Cirelli and Tononi, 2000).

However, there may be more heterogeneity in responses to SD across the brain than previously thought. For example, SD-driven transcript changes may vary between different brain structures (Mackiewicz et al., 2007; Terao et al., 2006; Vecsey et al., 2012). We have recently shown that while SD increases expression of the plasticity-mediating IEG *Arc* and Arc protein abundance in neocortical areas (e.g., primary somatosensory cortex; S1), it simultaneously decreases *de novo* synthesis of Arc in the hippocampal dentate gyrus (DG). Indeed, recent data have suggested that SD could differentially impact neuronal activity and dendritic spine density in hippocampal vs. neocortical structures (de Vivo et al., 2017; Havekes and Aton, 2020; Havekes et al., 2016; McDermott et al., 2003; Ognjanovski et al., 2018; Raven et al., 2019; Vyazovskiy et al., 2009). Because cognitive processes reliant on the hippocampus, such as episodic memory consolidation (Havekes and Abel, 2017; Saletin and Walker, 2012), seem particularly susceptible to disruption by SD, a critical unanswered question is whether SD differentially impacts network activity and plasticity in the two structures. Beyond this, within brain structures, there may be heterogeneity in the responses of different neuronal subtypes to SD. For example, within the neocortex, fast-spiking interneurons, or neurons with greater firing rates, appear to have differential firing rate changes across periods of sleep (Clawson et al., 2018; Vyazovskiy et al., 2009). Consistent with this idea, synaptic excitatory-inhibitory (E-I) balance was recently shown to vary in neocortex over the course of the day in a sleep-dependent manner (Bridi et al., 2020). Moreover, while most neocortical neurons fire at lower rates during slow wave sleep (SWS) vs. wake, some subclasses of neocortical neurons are selectively sleep-active (Gerashchenko et al., 2008).

Here we aimed to better characterize brain region- and cell type-specific changes evoked in the nervous system during SD. We used cell type-specific translating ribosome affinity purification (TRAP) (Sanz et al., 2019) to profile SD-mediated changes in ribosome-associated mRNAs in two prominent hippocampal and neocortical cell types – Camk2a+ pyramidal neurons and PV+ interneurons. Because interactions between these two cell types are critical for mediating state-dependent sensory plasticity and memory consolidation (Aton et al., 2013; Kuhlman et al., 2013; Ognjanovski et al., 2018; Ognjanovski et al., 2017), we characterized SD-driven changes in ribosome-associated transcripts encoding transcription-regulating IEGs, plasticity effector proteins, and circadian clock components in these two cell types. We find that SD generally causes more modest changes to these transcripts in hippocampal vs. neocortical circuits, and in PV+ interneurons vs. Camk2a+ neurons. Together our data suggest that the effects of SD on the brain are more heterogeneous than previously thought, and indicate region- and cell type-dependent differences in SD’s impact which may have important implications for brain function.

## Materials and Methods

### Mouse handling and husbandry

All animal procedures were approved by the University of Michigan Institutional Animal Care and Use Committee (PHS Animal Welfare Assurance number D16-00072 [A3114-01]). Animals were maintained on a 12:12h light/dark cycle (lights on at 8AM) with food and water provided *ad lib*. Mice expressing Cre recombinase in Camk2a+ neurons or PV+ interneurons (B6.Cg-Tg(Camk2a-cre)T29-1Stl/J or B6;129P2-*Pvalb*^*tm1(cre)Arbr*^/J; Jackson) were crossed to RiboTag mice (B6N.129-Rpl22^*tm1.1Psam*^/*J*; Jackson) to express HA-tagged Rpl22 protein in these neuron populations. 3-5 month old mice were individually housed one week prior to all experiments (with beneficial enrichment), and were habituated to handling for five days prior to experiments. Following habituation, and beginning at lights on (ZT0), mice were either allowed *ad lib* sleep in their home cage or were sleep deprived by gentle handling (Delorme et al., 2019; Durkin and Aton, 2016; Durkin et al., 2017). For sleeping animals, sleep behavior was visually scored at 5-min or 2-min intervals (for 6-h and 3-h SD, respectively), based on immobility and assumption of characteristic sleep postures. Previous research from our lab has shown that wake time over the final 45 min of the experiment correlates with *Arc* IEG expression in neocortex (Delorme et al., 2019). Thus to reduce the probability of confounding results from freely-sleeping mice, mice in the Sleep groups that spent > 60% of the final 45 min of the experiment in wake were excluded from subsequent analysis. All mice were sacrificed with an overdose of pentobarbital (Euthasol) prior to tissue harvest.

### Translating Ribosome Affinity Purification (TRAP)

TRAP was performed using methods described in prior studies (Sanz et al., 2009), with minor modifications. Following 3-6 h of *ad lib* sleep or SD, animals were euthanized with an overdose of pentobarbitol (Euthasol), their brains extracted, and hippocampi/cortices dissected in dissection buffer (1x HBSS, 2.5 mM HEPES [pH 7.4], 4 mM NaHCO_3_, 35 mM glucose, 100 μg/ml cycloheximide). Tissue was then transferred to glass dounce column containing 1 mL of homogenization buffer (10 mM HEPES [pH 7.4], 150 mM KCl, 10 mM MgCl_2_, 2 mM DTT, cOmplete™ Protease Inhibitor Cocktail [Sigma-Aldrich, 11836170001], 100 U/mL RNasin® Ribonuclease Inhibitors [Promega, N2111], and 100 μg/mL cycloheximide) and manually homogenized on ice. Homogenate was transferred to a 1.5 mL LoBind tubes (Eppendorf) and centrifuged at 1000×g at 4°C for 10 min. Supernatant was then transferred to a new tube, 90 μL of 10% NP40 was added, and samples were allowed to incubate for 5 min. Following this step, the supernatant was centrifuged at maximum speed for 10 min at 4°C, transferred to a new tube, and mixed with 10 μl of HA-antibody (Abcam, ab9110) (Jiang et al., 2015; Shigeoka et al., 2018). Antibody binding proceeded by incubating the homogenate-antibody solution for 1.5 hours at 4°C with constant rotation. During the antibody rinse, 200 μl of Protein G Dynabeads (ThermoFisher, 10009D) were washed 3 times in 0.15 M KCl IP buffer (10mM HEPES [pH 7.4], 150 mM KCl, 10 mM MgCl_2_, 1% NP-40) and incubated in supplemented homogenization buffer (10% NP-40). Following this step, supplemented buffer was removed, the homogenate-antibody solution was added directly to the Dynabeads, and the solution incubated for 1 h at 4°C with constant rotation. After incubation, the RNA-bound beads were washed four times in 900 μL of 0.35 M KCl (10 mM HEPES [pH 7.4], 350 mM KCl, 10 mM MgCl_2_, 1% NP40, 2 mM DTT, 100 U/mL RNasin® Ribonuclease Inhibitors [Promega, N2111], and 100 μg/mL cycloheximide). During the final wash, beads were placed onto the magnet and moved to room temperature. After removing the supernatant, RNA was eluted by vortexing the beads vigorously in 350 μl RLT (Qiagen, 79216). Eluted RNA was purified using RNeasy Micro kit (Qiagen).

### Quantitative real-time PCR (qPCR) and stability analysis

Quantitative real-time PCR (qPCR) experiments were performed as described previously (Delorme et al., 2019). Briefly, purified mRNA samples were quantified by spectrophotometry (Nanodrop Lite; ThermoFisher) and diluted to equal concentrations. 20-500 ng of mRNA was used to synthesize cDNA using iScript’s cDNA Synthesis Kit (Bio-Rad), cDNA diluted 1:10 in RNAse-free H_2_O, and measured using a CFX96 Real-Time System. Primers were designed for these studies, with the exception of Homer1a, for which sequences were established in a prior study (Mikhail et al., 2017). Primer specificity was confirmed using NIH Primer Blast (see **Table S1** for primer sequences). Three technical replicates were used for each sample. Relative changes in gene expression between sleep and SD were quantified using the ΔΔCT method, and these fold changes are presented on a log scale (log_2_ transformed value equivalent to ΔΔCT) with propagated errors. All statistical analyses were performed on ΔCT values.

Reference (housekeeping) genes for normalization were chosen for each experiment based on three measures: intragroup variability, intergroup variability, and an overall stability measure derived from total variance. Special emphasis was placed on selecting pairs of reference transcripts with countervailing intergroup differences. These measures were calculated using Normfinder (Andersen et al., 2004) and RefFinder (Xie et al., 2012) software. Because Normfinder is better suited for large sample sizes, RefFinder was used to validate Normfinder rankings and ensure genes with low (or opposite-direction) intergroup variability were chosen as housekeeping pairs. Stability measures were calculated for each sleeping condition, region, and mRNA population and repeated for mRNAs purified from *PV::RiboTag* and *Camk2a::Ribotag* mice (**Table 1**). The arithmetic mean of each housekeeping pair was then used to normalize target gene expression. As a final measure of housekeeping stability, we calculated each pairs’ fold change between mice in SD and Sleep groups.

**Table 1.** Housekeeping pairs used for RiboTag conditions. Change in gene expression presented as ratio^1^ and fold change^2^.

| mRNA Population | Condition | Region | Gene Pair | $SD(2^{-CT})/S(2^{-CT})^1$ | Fold Change<sup>2</sup> |
| --- | --- | --- | --- | --- | --- |
| Camk2a | 3hrs | Cortex | *Actg1/Hprt1* | 0.98 | -1.02 |
|  |  | Hippocampus | *Gapdh/Tuba4a* | 0.90 | -1.11 |
|  | 6hrs | Cortex | *Pgk1/Tbp* | 0.87 | -1.15 |
|  |  | Hippocampus | *Gapdh/Tuba4a* | 0.92 | -1.08 |
| Parvalbumin | 3hrs | Cortex | *Actg1/Hprt1* | 0.82 | -1.22 |
|  |  | Hippocampus | *Gapdh/Tuba4a* | 1.02 | 1.02 |
|  | 6hrs | Cortex | *Pgk1/Tbp* | 0.97 | -1.03 |
|  |  | Hippocampus | *Gapdh/Tuba4a* | 1.02 | 1.02 |

### RNAScope *in situ* hybridization

Fluorescent *in situ* hybridization was performed on 14-μm coronal sections taken from fixed-frozen brains of Sleep (*n* = 6) and SD (*n* = 6) mice. Section coordinates (1–3.0 mm lateral, −1.4 to −2.8 mm posterior to Bregma) were similarly distributed between Sleep and SD conditions (**Figure S2C**).The RNAScope Multiplex Fluorescent Reagent Kit v2 with 4-plex ancillary kit was used to label *Arc, Homer1a, Cfos,* and *Pvalb* transcripts (**Figure S2**). Prior to probe incubation, slices were pretreated with hydrogen peroxide (10 min, room temperature), Target Retrieval Reagent (5 min 99°C), and RNscope Protease III (30 min, 40°C). Slices were incubated with custom-synthesized *Arc* (20 bp, Target Region: 23-1066, 316911-C3, Advanced Cell Diagnostics), *Cfos* (20 bp, Target Region: 407-1427, 316921-C1, Advanced Cell Diagnostics), *Homer1a* (6 bp, Target Region: 1301-1887m 433941-C2, Advanced Cell Diagnostics), and *Pvalb* 16 (16 bp, Target RegionL 2-885, 421931-C4, Advanced Cell Diagnostics). Probes were chosen so as to overlap with regions amplified by qPCR primer pairs (**Table 2**). *Arc, Cfos, Homer1a, and Pvalb* were hybridized to Opal Dyes 620 (FP1495001KT, Akoya Biosciences), 570 (FP1488001KT, Akoya Biosciences), 690 (FP1497001KT, Akoya Biosciences), and 520 (FP1487001, Akoya Biosciences), respectively, for visualization. Positive and negative control probes were used in parallel experiments to confirm specificity of hybridization.

**Table 2.**
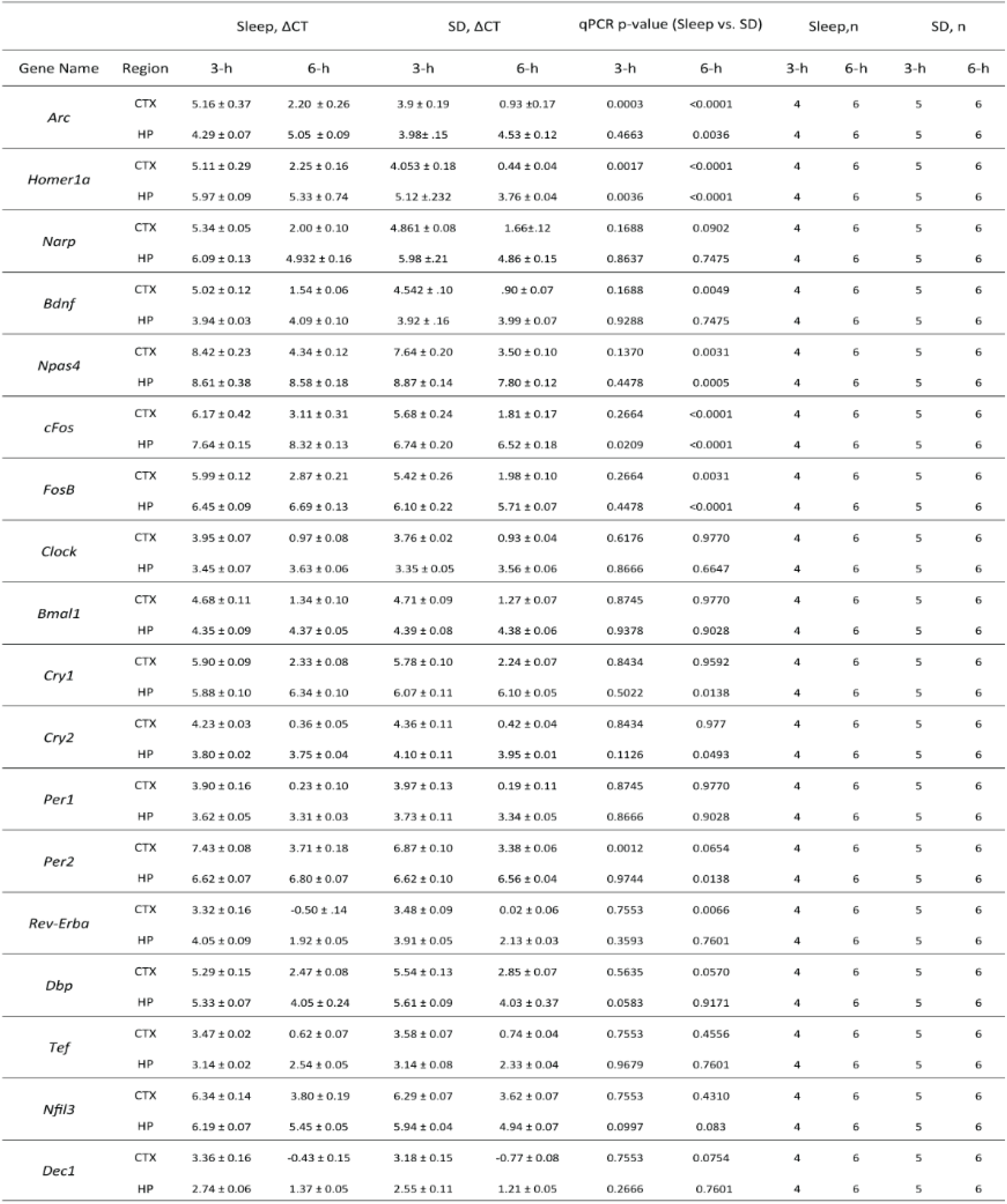
Sleep induced changes in ribosome-associated transcript abundance within Camk2a:RiboTag mice.

**Table 3.** Sleep induced changes in ribosome-associated transcript abundance within PV:RiboTag mice.

| Gene Name | Region | Sleep, ΔCT | | SD, ΔCT | | qPCR p-value (Sleep vs. SD) | | Sleep, n | | SD, n | |
| --- | --- | --- | --- | --- | --- | --- | --- | --- | --- | --- | --- |
|  |  | 3-h | 6-h | 3-h | 6-h | 3-h | 6-h | 3-h | 6-h | 3-h | 6-h |
| *Arc* | CTX | 6.64 ± 0.23 | 6.87 ± 0.07 | 5.25 ± 0.43 | 5.40 ± 0.07 | 0.0002 | <0.0001 | 4 | 5 | 4 | 6 |
| | HP | 6.67 ± 0.09 | 5.46 ± 0.14 | 6.58 ± 0.11 | 5.29 ± .19 | 0.9758 | 0.8837 | 4 | 6 | 5 | 6 |
| *Homer1a* | CTX | 7.49 ± 0.08 | 6.71 ± 0.13 | 7.03 ± 0.10 | 5.27 ± 0.12 | 0.3025 | <0.0001 | 4 | 5 | 4 | 6 |
| | HP | 9.02 ± 0.23 | 7.32 ± 0.35 | 9.00 ± 0.23 | 7.04 ± 0.27 | 0.9758 | 0.8673 | 4 | 6 | 5 | 6 |
| *Narp* | CTX | 8.22 ± 0.08 | 7.39 ± 0.07 | 7.77 ± 0.13 | 7.01 ± 0.09 | 0.3025 | 0.0196 | 4 | 5 | 4 | 6 |
| | HP | 8.60 ± 0.15 | 8.58 ± 0.47 | 8.71 ± 0.19 | 8.22 ± 0.21 | 0.9758 | 0.8383 | 4 | 6 | 5 | 6 |
| *Bdnf* | CTX | 8.03 ± 0.12 | 7.36 ± 0.19 | 7.89 ± 0.14 | 6.79 ± 0.09 | 0.6257 | 0.0014 | 4 | 5 | 4 | 6 |
| | HP | 0.60 ± 0.21 | 5.38 ± 0.26 | 1.17 ± 0.27 | 5.44 ± 0.20 | 0.2158 | 0.8907 | 4 | 6 | 5 | 6 |
| *Npas4* | CTX | 9.14 ± 0.43 | 7.97 ± 0.13 | 7.96 ± 0.14 | 7.11 ± 0.10 | 0.0143 | 0.0008 | 4 | 5 | 4 | 6 |
| | HP | 9.15 ± 0.20 | 5.21 ± 0.31 | 9.36 ± 0.13 | 5.40 ± 0.12 | 0.2737 | 0.6749 | 4 | 6 | 5 | 6 |
| *cFos* | CTX | 7.81 ± 0.10 | 7.04 ± 0.13 | 6.85 ± 0.38 | 5.21 ± 0.13 | 0.0336 | <0.0001 | 4 | 5 | 4 | 6 |
| | HP | 9.81 ± 0.13 | 7.56 ± 0.32 | 9.51 ± 0.10 | 5.98 ± 0.18 | 0.2081 | 0.0042 | 4 | 6 | 5 | 6 |
| *FosB* | CTX | 7.93 ± 0.16 | 7.05 ± 0.17 | 7.68 ± 0.15 | 6.22 ± 0.20 | 0.5072 | 0.0008 | 4 | 5 | 4 | 6 |
| | HP | 10.86 ± 0.12 | 6.65 ± 0.45 | 10.42 ± 0.11 | 7.10 ± 0.40 | 0.0814 | 0.5437 | 4 | 6 | 5 | 6 |
| *Clock* | CTX | 3.58 ± 0.07 | 3.29 ± 0.09 | 3.47 ± 0.04 | 3.04 ± 0.06 | 0.8513 | 0.2076 | 4 | 5 | 4 | 6 |
| | HP | 4.11 ± 0.05 | 4.06 ± 0.58 | 4.02 ± 0.05 | 2.68 ± 0.08 | 0.9716 | 0.9770 | 4 | 6 | 5 | 6 |
| *Bmal1* | CTX | 5.42 ± 0.09 | 4.51 ± 0.08 | 5.25 ± 0.07 | 4.51 ± 0.04 | 0.6893 | 0.9848 | 4 | 5 | 4 | 6 |
| | HP | 5.86 ± 0.09 | 3.71 ± 0.56 | 5.93 ± 0.09 | 5.59 ± 0.12 | 0.9716 | 0.9770 | 4 | 6 | 5 | 6 |
| *Cry1* | CTX | 8.57 ± 0.08 | 4.45 ± 0.03 | 8.63 ± 0.13 | 4.51 ± 0.05 | 0.8942 | 0.9436 | 4 | 5 | 4 | 6 |
| | HP | 6.24 ± 0.04 | 3.98 ± 0.60 | 6.15 ± 0.07 | 4.22 ± 0.23 | 0.9716 | 0.9099 | 4 | 6 | 5 | 6 |
| *Cry2* | CTX | 6.37 ± 0.05 | 3.12 ± 0.07 | 6.42 ± 0.09 | 3.09 ± 0.06 | 0.8942 | 0.979 | 4 | 5 | 4 | 6 |
| | HP | 5.18 ± 0.05 | 3.63 ± 0.61 | 5.09 ± 0.04 | 2.82 ± 0.07 | 0.9716 | 0.9994 | 4 | 6 | 5 | 6 |
| *Per1* | CTX | 4.16 ± 0.12 | 2.61 ± 0.10 | 3.92 ± 0.09 | 2.45 ± 0.04 | 0.4605 | 0.5214 | 4 | 5 | 4 | 6 |
| | HP | 4.66 ± 0.11 | 3.57 ± 0.53 | 4.56 ± 0.05 | 2.17 ± 0.08 | 0.9716 | 0.9994 | 4 | 6 | 5 | 6 |
| *Per2* | CTX | 7.70 ± 0.23 | 6.34 ± 0.15 | 7.21 ± 0.03 | 5.95 ± 0.15 | 0.0121 | 0.0113 | 4 | 5 | 4 | 6 |
| | HP | 6.98 ± 0.23 | 4.46 ± 0.89 | 6.88 ± 0.11 | 5.05 ± 0.24 | 0.9716 | 0.6197 | 4 | 6 | 5 | 6 |
| *Rev-Erba* | CTX | 3.89 ± 0.03 | 2.51 ± 0.06 | 3.91 ± 0.02 | 2.47 ± 0.07 | 0.9536 | 0.9276 | 4 | 5 | 4 | 6 |
| | HP | 4.97 ± 0.07 | 2.34 ± 0.09 | 4.96 ± 0.06 | 2.26 ± 0.11 | 0.9993 | 0.9805 | 4 | 6 | 5 | 6 |
| *Dbp* | CTX | 6.42 ± 0.07 | 5.47 ± 0.10 | 6.35 ± 0.09 | 5.49 ± 0.07 | 0.7058 | 0.9276 | 4 | 5 | 4 | 6 |
| | HP | 7.62 ± 0.08 | 6.12 ± 0.23 | 7.63 ± 0.10 | 6.10 ± 0.25 | 0.9993 | 0.9805 | 4 | 6 | 5 | 6 |
| *Tef* | CTX | 3.98 ± 0.02 | 3.35 ± 0.06 | 3.96 ± 0.02 | 3.24 ± 0.05 | 0.9536 | 0.7245 | 4 | 5 | 4 | 6 |
| | HP | 4.80 ± 0.09 | 3.23 ± 0.05 | 4.80 ± 0.02 | 2.94 ± 0.07 | 0.9993 | 0.7872 | 4 | 6 | 5 | 6 |
| *Nfil3* | CTX | 7.11 ± 0.05 | 6.46 ± 0.17 | 6.88 ± 0.06 | 6.13 ± 0.07 | 0.0102 | 0.0206 | 4 | 5 | 4 | 6 |
| | HP | 7.86 ± 0.10 | 8.58 ± 0.47 | 7.71 ± 0.13 | 8.22 ± 0.21 | 0.6950 | 0.713 | 4 | 6 | 5 | 6 |
| *Dec1* | CTX | 4.88 ± 0.04 | 3.62 ± 0.09 | 4.66 ± 0.02 | 3.27 ± 0.04 | 0.0102 | 0.0143 | 4 | 5 | 4 | 6 |
| | HP | 5.54 ± 0.11 | 3.92 ± 0.11 | 5.49 ± 0.07 | 3.82 ± 0.07 | 0.9951 | 0.9805 | 4 | 6 | 5 | 6 |

### Immunohistochemistry

For immunohistochemical validation of appropriately cell targeted HA expression in RiboTag-expressing mice, *Camk2a::RiboTag* and *PV::RiboTag* mice from Sleep (*n* = 6) and SD (*n* = 6) groups were sacrificed and perfused with PBS followed by 4% paraformaldehyde. 50-μm brain sections were blocked with normal goat serum for 2 h and incubated overnight using biotin-conjugated anti-HA (Biolegend 901505, 1:500) and anti-parvalbumin (Synaptic Systems 195 004, 1:500) antibodies at 4°C. The following day, sections were stained with Streptavidin-Alexa Fluor® 647 (Biolegend 405237) and Alexa Fluor® 555 Goat Anti-Guinea pig IgG H&L (Abcam ab150186). Stained sections were coverslipped in ProLong Gold Antifade Reagent (ThermoFisher, P36930). Fluorescence intensity was used to identify HA-expressing (HA^+^) cells, PV-expressing (PV^+^) cells, and overlapping cells within the DG, CA1, CA3, and neocortex. To account for differences in localization and spread of antibody staining, both PV+ HA-expressing cells and HA^+^ PV-expressing cells were identified, and overlap was quantified in terms of both cell count and cell area (**Figure 1**). Quantification was performed using the semi-automated protocol detailed below.

**Figure 1.**
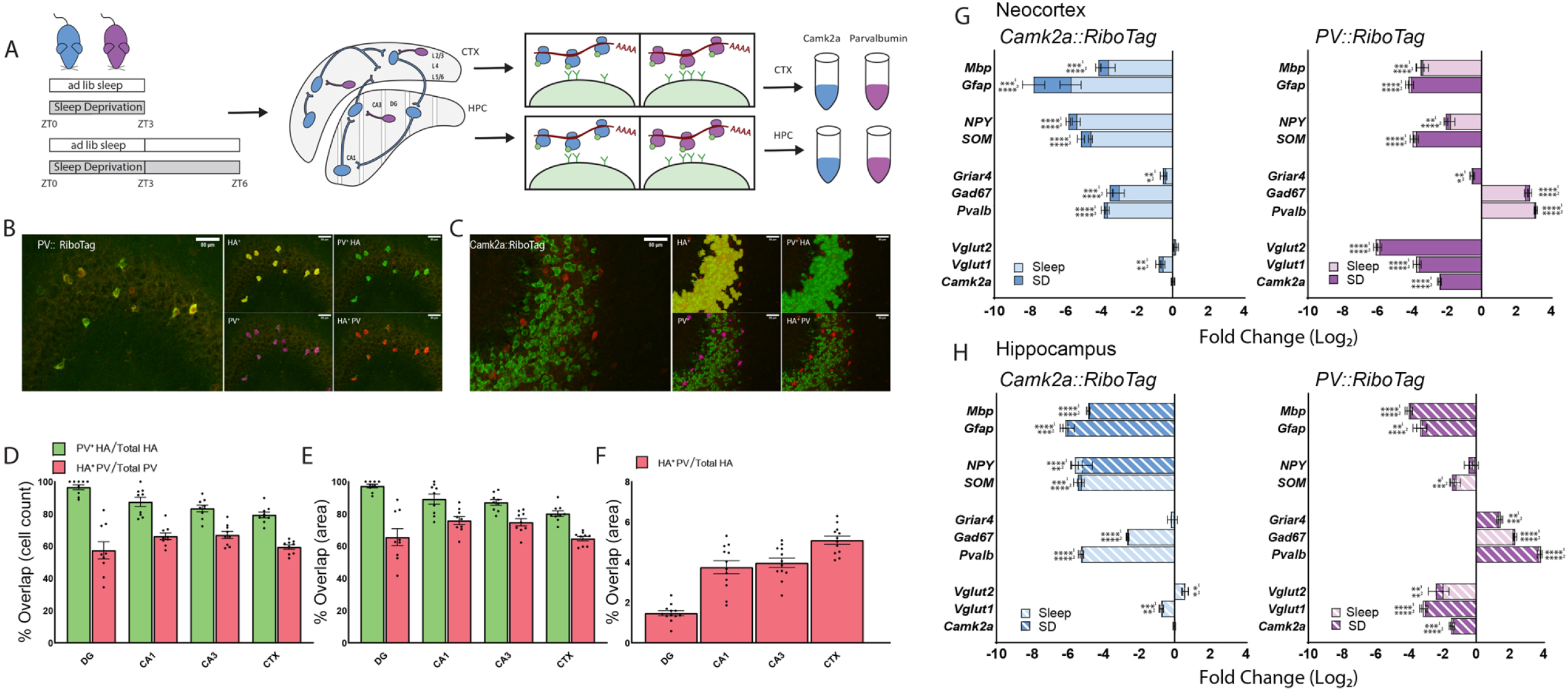
Experimental design and validation of TRAP. (**A**) *Camk2a::RiboTag* (blue) and *PV::RiboTag* (magenta) transgenic mice were sacrificed after a 3- or 6-h period of *ad lib* sleep (Sleep) or sleep deprivation (SD) starting at lights on (ZT0). Ribosome-associated mRNAs were affinity purified from hippocampus and neocortex. (**B**) HA expression in PV+ interneurons was validated with immunohistochemistry in *PV::RiboTag* mice, with automated detection of HA (green fluorescence, labeled in yellow) and PV (red fluorescence, labeled in purple) expression. Areas of overlapping fluorescence were then identified; to account for differences in antibody staining, both PV^+^ HA-expressing areas (green) and HA^+^ PV-expressing areas (red) were identified. Scale bars = 50 μm. (**C**) Example of automated protocol used in *Camk2a::RiboTag* mice to quantify non-specific expression. (**D-E**) HA expression presented as proportion of overlapping cells vs. total cell count (**D**) and total area (**E**) in *PV::RiboTag* sections. (**F**) PV^+^ HA-expressing areas over total HA^+^ area in *Camk2a::RiboTag* sections. (**G**) Enrichment of markers for glia (*Mbp, Gfap*), non-PV+ inhibitory neurons (*NPY, SOM*), PV+ neurons (*Griar4, Gad67, PV*), and excitatory neurons (*Vglut1, Vglut2, Camk2a*) calculated as ΔΔCT between affinity purified (RiboTag) mRNA and Input mRNA from neocortex. Data presented as log(2) transformed fold changes. (**H**) Enrichment values for *Camk2a::RiboTag* and *PV::RiboTag* hippocampi. Gene expression was normalized to housekeeping gene pairs according to their respective condition (see **Table 1**). Values indicate mean ± SEM with propagated error; *, **, ***, and **** indicate *p* < 0.05, *p* < 0.01, *p* < 0.001, and *p* < 0.0001, respectively, one sample t-test against hypothetical value of 0.

### Imaging and quantification

RNAScope probe fluorescence signals were captured and stitched using a 40× objective lens on a Leica 3D STED SP8. Immunostained brain sections were imaged on a Leica SP5 laser scanning confocal microscope. Settings were fixed for each imaging session. Fluorescence images were analyzed using MIPAR image analysis software in their raw grayscale format (Sosa et al., 2014). Two images per region (one per hemisphere) were quantified for each animal. Total fluorescence dot number and average intensity of a single dot calculated as described here (ACDBio, 2017), for PV+ and non-PV+ regions of interest (ROIs) within granule (dentate gyrus), pyramidal (CA1, CA3), and cortical layers 1-6 (layers were manually isolated using a freehand tool by a scorer blind to experimental condition). Fluorescence intensity and expression overlap were calculated using a semi-automated protocol curated by blinded scorer. Briefly, a non-local means filter was used to reduce image noise, and an adaptive threshold was used to identify areas > 30 μm whose mean pixel intensity was 200% of its surroundings. Identified areas were labeled as IEG or PV and manually edited to refine labeling, select for representative dots, and remove artifacts (manual editing was not used to label any additional IEG expression). Finalized labeling was used to delineate PV+ and non-PV+ ROIs, select for background area (area in the ROI minus areas of labeled expression), and identify IEG+ PV+ cells (**Figure S3**). Intensity values from ROIs, background, and selected dots were used to calculate fluorescence dots/area. Average background intensity was calculated as the fluorescence intensity of the selected background area per unit area. The average intensity of a single fluorescent dot was calculated for each transcript as the intensity of manually selected representative dots within the ROI, minus the average background intensity multiplied by the area, divided by the total number of selected dots. Dot intensity values did not differ between Sleep and SD mice for specific transcripts. The total fluorescent dot number within each ROI was calculated by subtracting average background intensity from total ROI fluorescence intensity, multiplied by total area, divided by average dot intensity.

## Results

### TRAP-based characterization of ribosome-associated transcripts in Camk2a+ and PV+ neuronal populations

To quantify how ribosome-associated transcripts in different neuronal populations within the hippocampus and neocortex are affected by sleep loss, we crossed RiboTag transgenic mice (with Cre recombinase-dependent expression of HA-tagged Rpl22 protein) to either Camk2a-Cre or PV-Cre transgenic lines (Sanz et al., 2019) (**Figure 1A**). Appropriate cell type-specific expression of Rpl22^HA^ in *Camk2a::RiboTag* and *PV::RiboTag* mice was verified using immunohistochemistry (**Figure 1B-F**). HA expression was largely circumscribed to the intended cell type. For example, 86.7 ± 1.5% and 79.4 ± 1.8% of HA+ neurons within the hippocampus and neocortex of *PV::RiboTag* mice co-expressed PV peptide. In both *Camk2a::RiboTag* and *PV:RiboTag* mice, expression of HA in non-targeted cell types of the hippocampus (based on lack of co-immunostaining for Camk2a or PV) was minimal (3.6 ± 0.2% and 13.3 ± 1.5%, respectively; **Figure 1D-F**).

We next validated cell type-specificity of ribosome-associated transcripts isolated from transgenic mouse lines. Following a period of *ad lib* sleep of sleep deprivation (SD) starting at lights on (ZT0), hippocampi and neocortex were dissected, and ribosome-associated mRNAs were isolated (Sanz et al., 2019). We compared abundance of cell type-specific transcripts between RiboTag affinity purified mRNA and Input mRNA from whole hippocampus or neocortex homogenate using qPCR. Enrichment or de-enrichment of these cell markers was compared with a null hypothetical value of 0 using one-sample t-tests. We found that ribosomal-associated transcripts from both the neocortex and hippocampus of *Camk2a::RiboTag* mice de-enriched for markers of glial cell types (*Mbp, Gfap*), non-PV+ inhibitory neurons (*Npy, Sst*), PV interneurons (*Gad67, Pvalb*), and *Vglut1* relative to Input (**Figure 1G-H**). Hippocampal enrichment patterns mirrored those of the neocortex with the exception of *Vglut2*, which was significantly enriched relative to Input. Ribosome-associated transcripts from *PV::RiboTag* mice de-enriched for markers of glial (*Mbp, Gfap*), non-PV+ inhibitory (*Npy, Sst*), and excitatory neurons (*Vglut1*, *Vglut2*, *Camk2a*) while enriching for PV+ interneuron markers (*Pvalb, Gad67*) relative to Input. We made comparisons of cell type-specific transcript enrichment separately for mice which were either allowed *ad lib* sleep or sleep deprived (SD) over the first 3 or 6 h after lights on (i.e., from ZT0-3, or ZT0-6; **Figure S1A**). No substantial differences in enrichment patterns were observed between Sleep and SD mice (*N.S.*, Holm-Sidak *post hoc* test). These data confirm the high degree of specificity of TRAP-based profiling for ribosomal transcripts from Camk2a+ principal neurons and PV+ interneurons.

### SD-driven changes in ribosome-associated plasticity-related mRNAs vary with cell type and brain structure

We first quantified a subset of transcripts encoding for proteins involved in synaptic plasticity (i.e., plasticity effectors) whose expression levels have been reported previously as altered by SD - *Arc, Homer1a, Narp, and Bdnf* (Cirelli et al., 2004; Maret et al., 2008). Ribosome-associated transcript abundance was first quantified in Camk2a+ neocortical and hippocampal neuron populations after 3 h of *ad lib* sleep (Sleep; *n* = 4) or SD (*n* = 5), starting at lights on (ZT0). Consistent with previous findings (Cirelli et al., 2004), 3-h SD significantly increased neocortical *Arc* (*p* < 0.001, Holm–Sidak *post hoc* test) and *Homer1a* (*p* < 0.01) (Maret et al., 2008) ribosome-associated mRNA (**Figure 2A**). In contrast, and consistent with recent data (Delorme et al., 2019), 3-h SD significantly increased *Homer1a* abundance on hippocampal ribosomes (*p* < 0.01), but did not significantly affect *Arc* abundance (*N.S*., Holm–Sidak *post hoc* test). Overall patterns of transcript abundance for the plasticity-regulating proteins *Bdnf* and *Narp* followed a similar trend, with unchanged levels in hippocampal Camk2a+ neurons (*N.S*, Holm–Sidak *post hoc* test), and modestly (but not significantly) increased levels in neocortical neurons (*Narp* and *Bdnf*, *N.S*.). After more prolonged (6-h) SD (*n* = 6 mice/group), ribosome-associated *Arc* (*p* < 0.0001), *Homer1a* (*p* < 0.0001), and *Bdnf* (*p* < 0.01) transcripts were all increased in neocortical Camk2a+ neurons, whereas *Arc* (*p* < 0.01) and *Homer1a* (*p* < 0.0001) were increased in hippocampal Camk2a+ neurons (**Figure 2B**).

**Figure 2.**
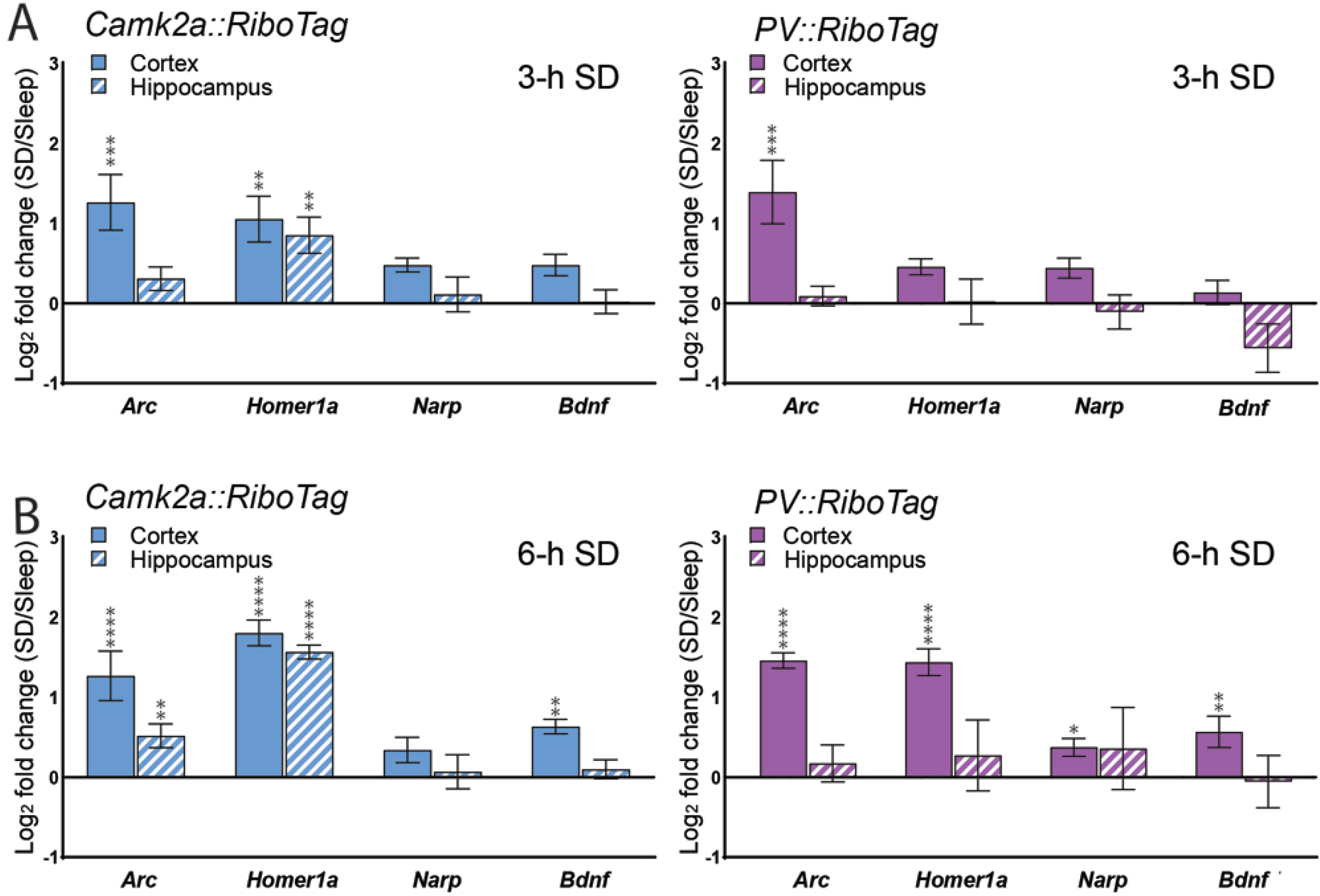
SD increases ribosome-associated plasticity effector transcripts in a cell type- and region-specific manner. **(A)** 3-h SD significantly increased *Arc* and Homer1alevels on ribosomes from Camk2a+ neocortical (solid) neurons; only *Homer1a* increased in hippocampal (dashed) neurons. 3-h SD significantly increased *Arc* on ribosomes from PV+ interneurons in neocortex; no significant change was observed in the hippocampal PV+ interneuron population. **(B)** *Arc*, *Homer1a*, and *Bdnf* significantly increased after 6-h SD in Camk2a+ neocortical neurons; *Arc* and *Homer1a* were increased within the Camk2a+ hippocampal population. All effector transcript levels were significantly elevated after 6-h SD in PV+ interneurons in neocortex; no significant change was observed in the hippocampal PV+ population. Transcript level changes are presented as a log_2_ fold change between SD and *ad lib* sleep mice. Values indicate mean ± SEM with propagated error; *, **, ***, and **** indicate *p* < 0.05, *p* < 0.01, *p* < 0.001, and *p* < 0.0001, respectively, Holm–Sidak *post hoc* test vs. Sleep.

We next quantified ribosome-associated transcript abundance in PV+ interneuron populations from the neocortex (*n* = 4 mice/group) and hippocampus (*n* = 4 and *n* = 5 mice for Sleep and SD). 3-h SD significantly increased *Arc* (*p* < 0.001, Holm–Sidak *post hoc* test) abundance in neocortical PV+ interneurons, but had no effect on transcript abundance for plasticity-related proteins in hippocampal PV+ interneurons (*N.S.*, Holm–Sidak *post hoc* test). 6-h SD increased abundance of these transcripts in the neocortical PV+ interneuron population (*n* = 5 and *n* = 6 mice for Sleep and SD) in a manner similar to the Camk2a+ neuronal population (*Arc*, *p* < 0.0001; *Homer1a*, *p* < 0.0001; *Narp*, *p* < 0.05; *Bdnf*, *p* < 0.01). In contrast, 6-h SD caused no significant change in any of the ribosome-associated transcripts’ abundance in hippocampal PV+ interneurons (*n* = 6 mice/group).

### SD differentially affects abundance of ribosome-associated mRNAs encoding activity-dependent transcription regulators based on cell type in hippocampus vs. neocortex

To better characterize how SD affects activity-regulated pathways in Camk2a+ and PV+ populations, we quantified ribosome-associated transcript abundance for IEGs encoding transcription regulatory factors - *Npas4*, *Cfos*, and *Fosb*. We first quantified transcript abundance in Camk2a+ neocortical and hippocampal neuronal populations after 3-h of *ad lib* sleep (Sleep; *n* = 4) or SD (*n* = 5), starting at lights on (ZT0). 3-h SD produced no significant change in ribosome-associated transcript abundance in Camk2a+ neocortical cells (*N.S*. for all transcripts, Holm– Sidak *post hoc* test) while significantly increasing *Cfos* abundance in the hippocampus (*p* < 0.05; **Figure 3A**). After prolonged (6-h) SD (*n* = 6 mice/group; **Figure 3B**), neocortical *Npas4* (*p* < 0.01), *Cfos* (*p* < 0.0001) and *Fosb* (*p* < 0.01) abundance increased on ribosomes in Camk2a+ neurons. In the hippocampus, ribosome-associated *Npas4* (*p* < 0.001), *Cfos* (*p* < 0.0001), and *Fosb* (*p* < 0.0001) all increased in abundance in Camk2a+ neurons after 6-h SD.

**Figure 3.**
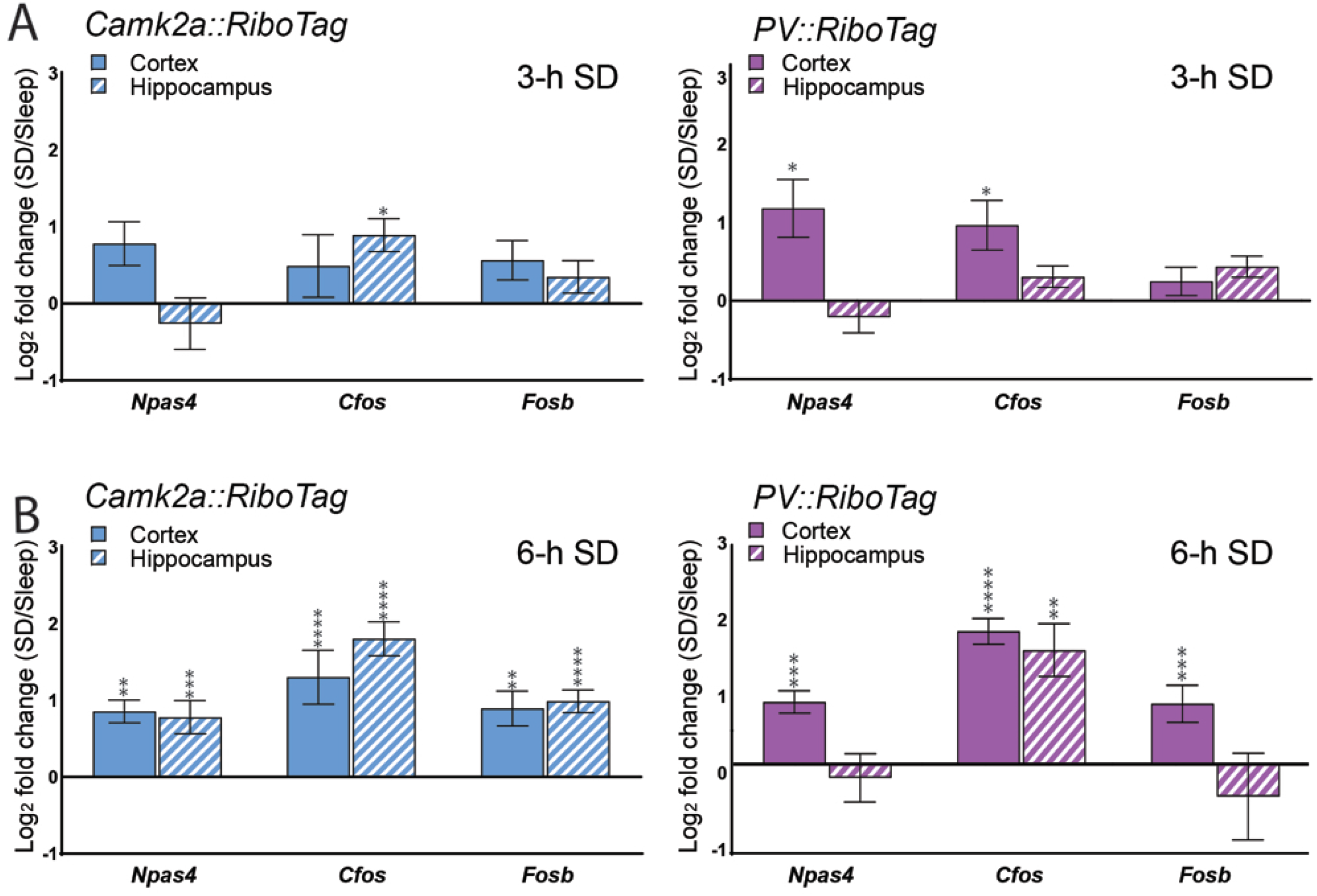
SD increases ribosome-associated transcripts encoding immediate-early transcription regulators in a cell type- and region-specific manner. **(A)** 3-h SD had no significant effect on IEG transcript levels on ribosomes from Camk2a+ neocortical (solid) neurons; only *Cfos* increased in hippocampal (dashed) neurons. 3-h SD significantly increased *Npas4* and *Cfos* on ribosomes from PV+ interneurons in neocortex, but did not affect IEG abundance on ribosomes from hippocampal PV+ neurons. **(B)** 6-h SD significantly increased *Npas4*, C*fos*, and *Fosb* levels in Camk2a+ neocortical neurons, Camk2a+ hippocampal neurons, and PV+ neocortical interneurons. Only *Cfos* significantly increased in the PV+ hippocampal population with 6-h SD. Transcript level changes are presented as a log_2_ fold change between SD and *ad lib* sleep mice. Values indicate mean ± SEM with propagated error; *, **, ***, and **** indicate *p* < 0.05, *p* < 0.01, *p* < 0.001, and *p* < 0.0001, respectively, Holm–Sidak *post hoc* test vs. Sleep.

We next quantified ribosome-associated transcripts encoding IEG transcription factors in PV+ interneurons from the neocortex (*n* = 4 mice/group) and hippocampus (*n* = 4 and *n* = 5 mice for Sleep and SD). 3-h SD significantly increased neocortical *Npas4* and *Cfos* (*p* < 0.05) abundance, but had no effect on transcript abundance in the hippocampus (*N.S.* for all transcripts, Holm–Sidak *post hoc* test). 6-h SD significantly increased all three transcripts’ abundance (*p* < 0.0001 for *Cfos, p* < 0.001 for all other transcripts) in the neocortex, but only affected *Cfos* in the hippocampus (*p* < 0.01). Overall, ribosome-associated transcript abundance in PV+ interneurons from the neocortex underwent fold changes that were slightly higher than hippocampus.

### Subregion- and layer-specific effects of SD on mRNA abundance in PV+ and non-PV+ neurons

Recent findings suggest that effects of SD on transcription and translation may be more region- and subregion-specific than previously thought (Delorme et al., 2019; Havekes and Aton, 2020). To more precisely characterize region-specific changes in mRNA abundance after SD, we used fluorescence *in situ* hybridization to visualize *Pvalb*, *Arc*, *Homer1a*, and *Cfos* transcripts in C57Bl6/J mice after 6-h SD (*n* = 6) or *ad lib* sleep (*n* = 5)(**Figure 4A, Figure 5A-B, Figure S1B, Figure S2**). Transcripts were quantified separately in neocortical layers 1-6 and DG, CA3, and CA1 hippocampal subregions. *Pvalb* expression was used to discriminate expression in PV+ interneurons from that in non-PV+ (mainly pyramidal) neurons. Regions of interest (ROIs) for PV+ interneurons and non-PV+ regions were identified separately and total transcript expression (total fluorescence dot number) was normalized to the area of their respective ROI. We first quantified mRNA abundance after Sleep vs. SD among non-PV+ cells in neocortical regions overlying dorsal hippocampus (including S1)(**Figure 4B**). Across neocortex as a whole, SD significantly increased *Arc* in non-PV+ neurons (Sleep = 24.8 ± 10.3 vs. SD = 79.2 ± 10.1 dots/mm^2^, *p* < 0.05, Holm– Sidak *post hoc* test), and showed a tendency for increasing *Cfos* (Sleep = 8.6 ± 3.9 vs. SD = 26.2 ± 5.1 dots/mm^2^, *p* = 0.053) and *Homer1a* (Sleep = 1.4 ± 0.5 vs. SD = 7.8 ± 2.6 dots/mm^2^, *p* = 0.056). Expression was also quantified in individual neocortical layers. The largest effects of SD were seen for *Homer1a* and *Cfos* in layers 4 (*Homer1a*: Sleep = 1.6 ± 0.6 vs. SD = 7.8 ± 2.2 dots/mm^2^, *Cfos*: Sleep = 13.5 ± 6.4 vs. SD = 40.5 ± 7.1 dots/mm^2^) and 5 (*Homer1a*: Sleep= 1.5 ± 0.4 vs SD=9.5 ± 2.8 dots/mm^2^, *Cfos*: Sleep = 8.8 ± 3.8 vs. SD = 34.5 ± 6.9 dots/mm^2^, *p* < 0.05). SD increased *Arc* dots/mm^2^ significantly across layers 2/3 (Sleep = 15.2 ± 5.8 vs. SD = 45.8 ± 3.7 dots/mm^2^, *p* < 0.01, unpaired *t*-test), 4 (Sleep = 36.3 ± 14.3 vs. SD=137.5 ± 17.7 dots/mm^2^, *p* < 0.01), and 5 (Sleep = 21.7 ± 8.2 vs. SD = 81.7 ± 12.8 dots/mm^2^, *p* < 0.05) (**Figure 4B**). No changes in expression were observed with SD in layer 6, and layer 1 expression was not analyzed due to low overall expression and cell density. In dramatic contrast to the relatively large changes in non-PV+ transcript abundance with SD in neocortex, neither *Arc* nor *Homer1a* (*N.S.*, Holm–Sidak *post hoc* test) levels were significantly altered by SD in any region of dorsal hippocampus (**Figure 5C**). *Cfos* was increased significantly with SD in CA3 only (Sleep = 2.8 ± 0.5 vs. SD = 10.7 ± 1.4 dots/mm^2^, *p* < 0.01) with no significant changes in CA1 or DG (*N.S.*, Student’s t-test). We then quantified transcript abundance within PV+ interneurons, using *Pvalb* mRNA expression to define the PV+ ROI (**Figure 4C**). Overall IEG expression in PV+ interneurons was relatively low. SD caused no significant changes in *Arc* or *Homer1a* in any layer of the neocortex, although *Cfos* dots/μm^2^ increased selectively in PV+ interneurons in layer 2/3 (Sleep = 0.014 ± 0.002 vs. SD = 0.043 ± .009 dots/μm^2^, *p* < 0.01). Because many PV+ interneurons expressed no detectable IEGs, we also quantified expression within the subpopulation of PV+ interneurons which had detectable levels of mRNA expression. Using a semi-automated protocol for this more circumscribed analysis, we found that SD did not affect expression levels for *Arc* or *Cfos*, but did increase *Homer1a* dots/μm^2^ when measured across the entire neocortex (**Figure 4C**). Consistent with limited ribosome-associated transcript changes in hippocampus with SD (**Figures 2** and **3**), no significant changes in IEG expression were observed in PV+ interneurons any region of dorsal hippocampus with SD, regardless of method for quantification (**Figure 5D-E**).

**Figure 4.**
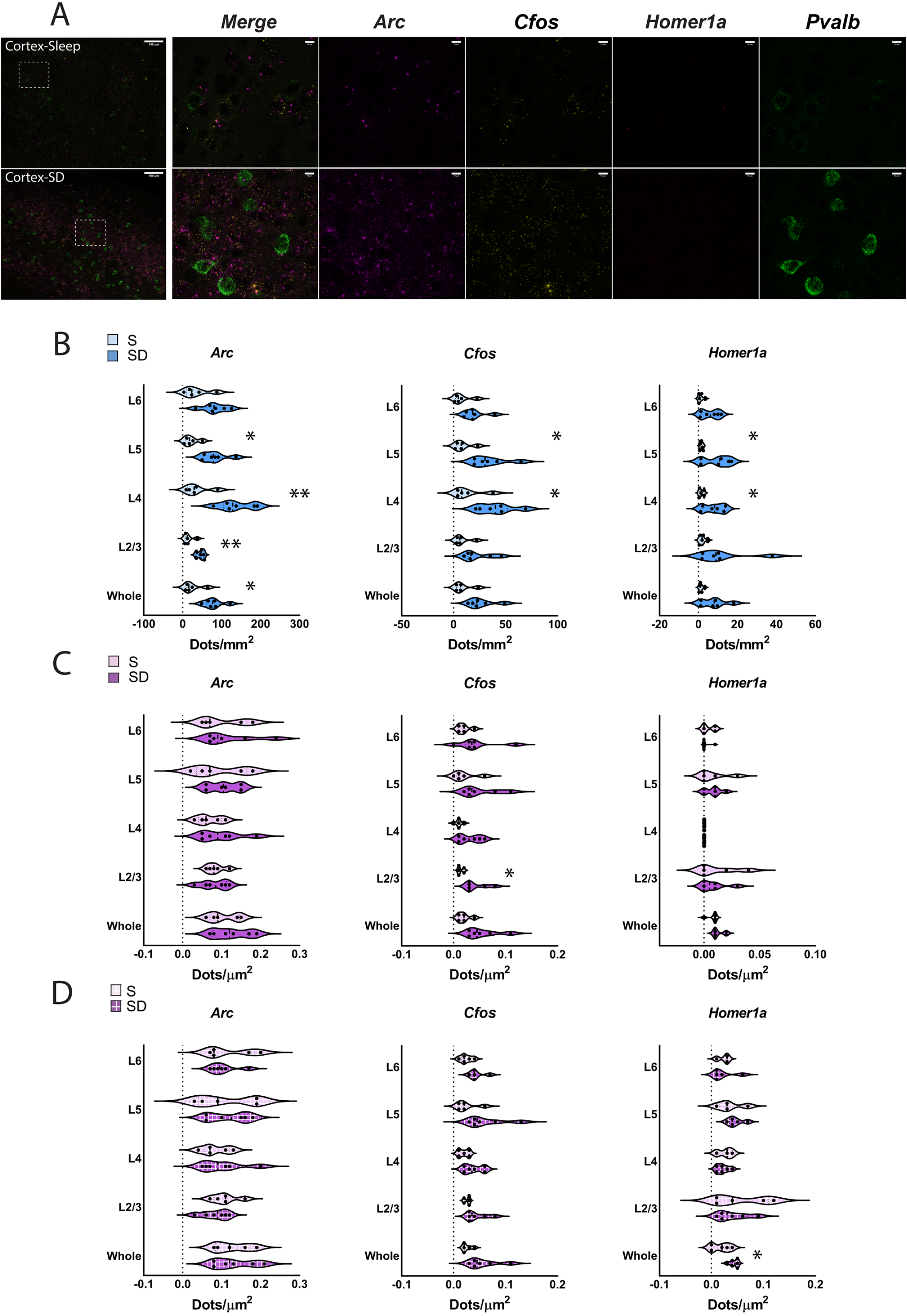
Layer-specific induction of IEG expression increases in neocortex after SD. **(A)** Representative images show neocortical IEG expression after 6 h of *ad lib* sleep (*n* = 5 mice) or SD (*n* = 6 mice). Inset regions are shown at higher magnification on right. Scale bars indicate 100 μm and 10 μm respectively. **(B)** 6-h SD significantly increased *Arc* dots/mm^2^ among non-PV+ cells in whole cortex and layers 2/3, 4 and 5, and *Cfos* and *Homer1a* dots/mm^2^ in layers 4 and 5. **(C)** 6-h SD significantly increased *Cfos* dots/μm^2^ among *Pvalb+* cells (magenta) in layer 2/3; no other significant changes were observed. **(D)** When analysis was restricted to IEG+ *Pvalb+* cells (magenta, box pattern), SD significantly increased *Homer1a* dots/μm^2^ among *Homer1a*+ *Pvalb+* cells in whole cortex; no other significant changes were observed. Violin plots show distribution of values for individual mice; * and ** indicates *p* < 0.05 and *p* < 0.01, Holm–Sidak *post hoc* test vs. Sleep.

**Figure 5.**
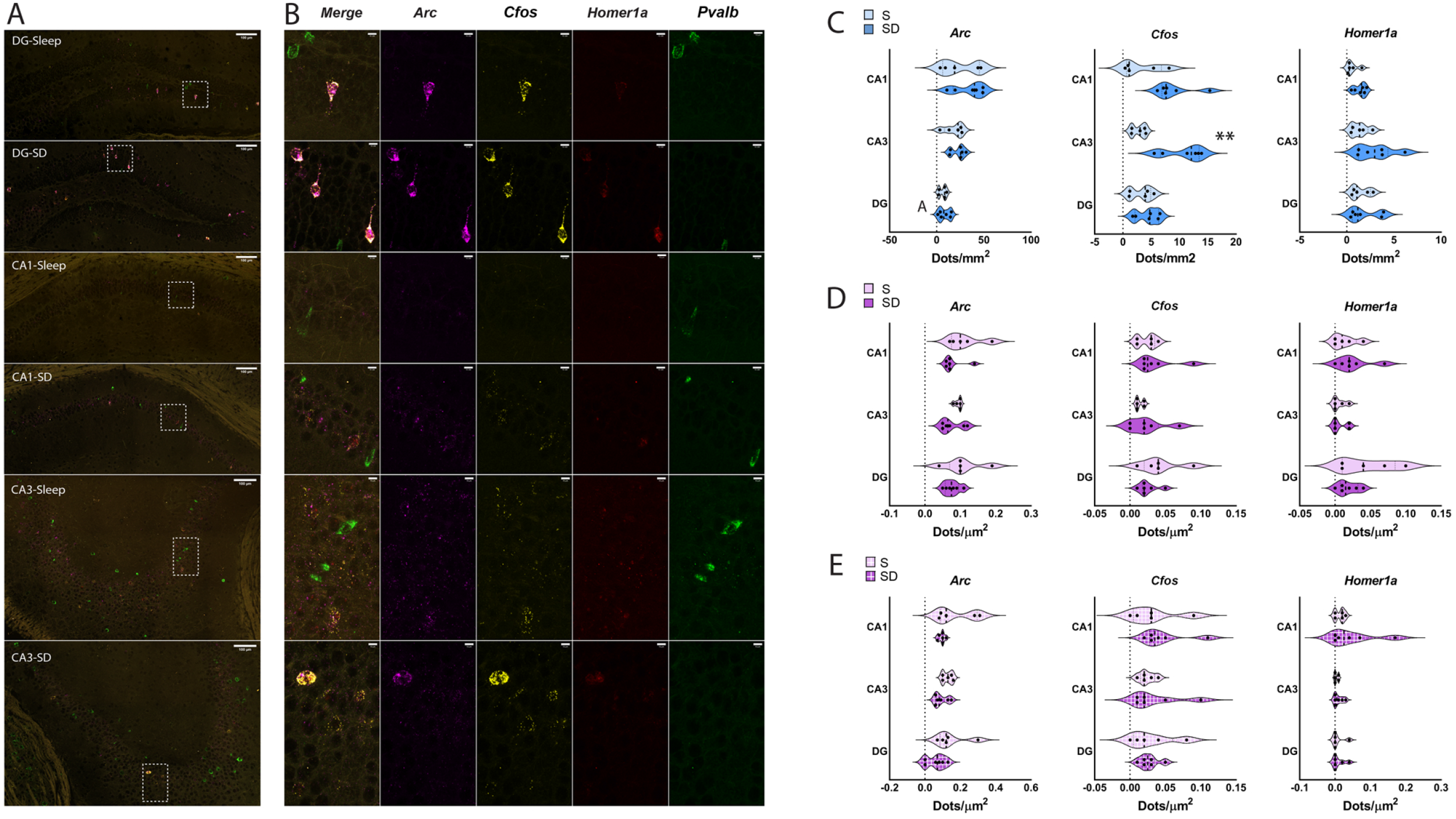
Cell type- and region-specific changes in hippocampal IEG expression after SD. **(A)** Representative images show IEG expression in DG, CA1, and CA3 after 6 h of *ad lib* sleep (*n* = 5 mice) or SD (*n* = 6 mice). Inset regions are shown at higher magnification on right. Scale bars indicate 100 μm and 10 μm respectively. **(B)** 6-h SD significantly increased *Cfos* dots/mm^2^ among non-PV+ (blue) cells in CA3; no other significant changes observed. **(C-D)** No significant changes were observed within DG, CA3, or CA1 in *Pvalb+* cells (magenta) **(C)** or IEG+ *Pvalb+* cells (magenta, box pattern) **(D)**. Violin plots show distribution of individual subjects; ** indicates *p* < 0.01, Holm–Sidak *post hoc* test vs. Sleep.

One possibility is that the relative proportion of IEG+ PV+ interneurons varied as a function of SD. Because PV+ interneurons varied substantially in terms of ROI size, we quantified the IEG+ proportion of PV+ interneurons in Sleep and SD mice, as a function of both cell count and ROI area (**Figure 6**). We found the SD significantly increased the proportion of *Arc*+ and *Cfos*+ PV+ interneurons in the neocortex, across all layers quantified (**Figure 6B**). No significant differences were observed in the proportion of *Homer1a*+ PV+ interneurons. Similarly, we found significant increases in the proportion of *Arc*+ and *Cfos*+ PV+ area after SD for all neocortical layers, with the exception of layer 5 (**Figure 6C**). No differences were observed for *Homer1a*+ area with PV+ interneurons using this measure. No significant changes in any of the mRNAs’ expression were observed after SD in PV+ interneurons in any region of the hippocampus after SD, regardless of the method of quantification (**Figure 6D-F**).

**Figure 6.**
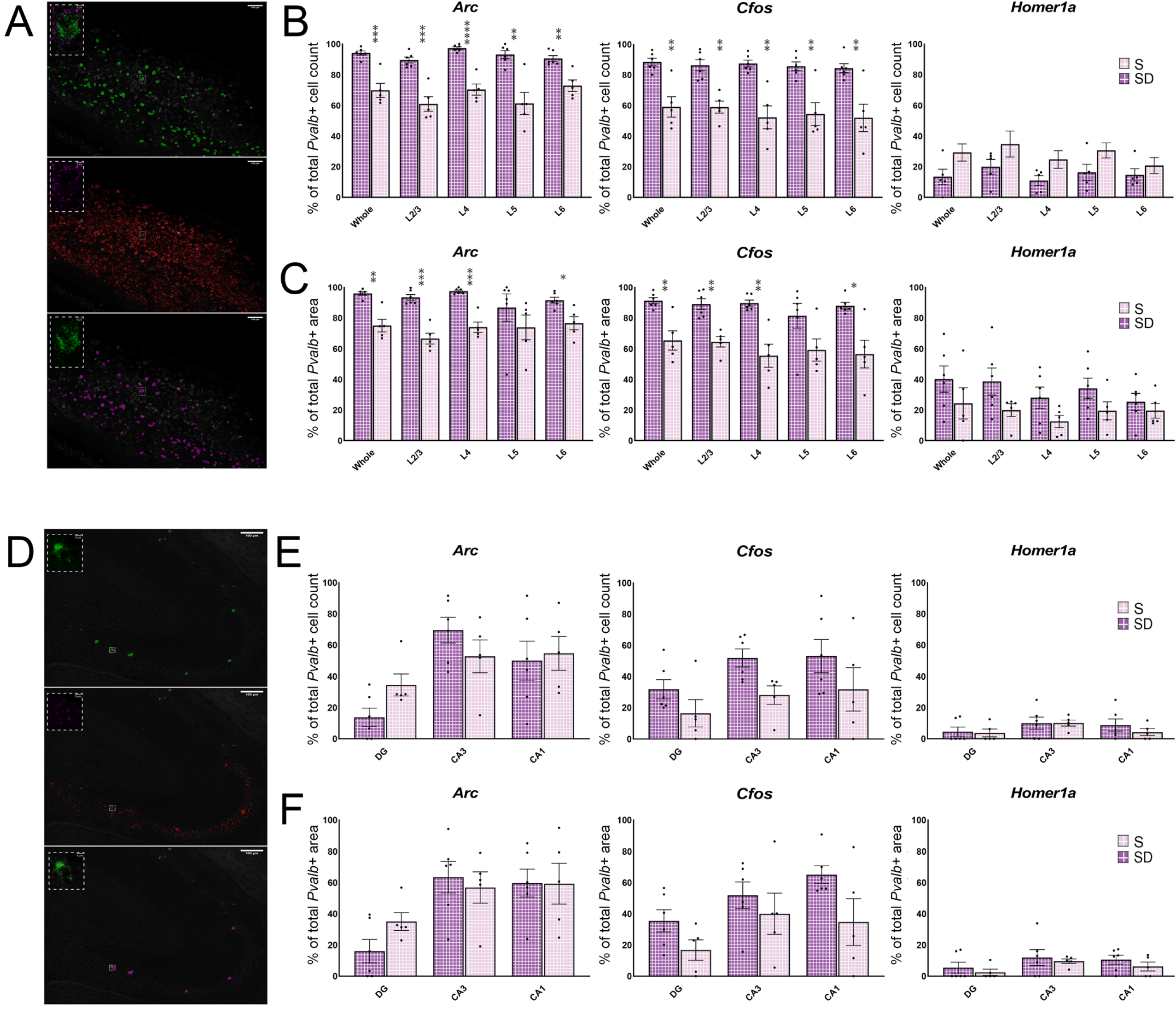
SD increases the proportion of IEG+ PV+ interneurons in neocortex, but not hippocampus. **(A)** An automated protocol identified *Pvalb* (green) and IEG (red) *in situ* fluorescence; cells with overlapping fluorescence were marked as IEG+ (magenta). Total IEG+ *Pvalb+* area was then calculated as the proportion of total *Pvalb+* area. **(B-C)** 6-h SD increased the proportion **(B)** and area **(C)** of *Pvalb+* cells expressing *Arc* or *Cfos*, but not *Homer1a*, across most neocortical layers. Values indicate mean ± SEM; *, **, ***, and **** indicate *p* < 0.05, *p* < 0.01, *p* < 0.001, and *p* < 0.0001, respectively, Holm–Sidak post hoc test vs sleep. **(D)** The same method identified IEG+ *Pvalb+* cells within hippocampal subregions DG, CA1, and CA3. **(E-F)** SD had no effect on the proportion **(E)** or area **(F)** of *Pvalb+* hippocampal cells expressing *Arc*, *Cfos*, or *Homer1a*, Values indicate mean ± SEM; *N.S.*, Holm–Sidak *post hoc* test vs. Sleep.

Critically, *Pvalb* expression itself can be regulated as a function of synaptic plasticity (Donato et al., 2013). We found that when expression values were calculated cell by cell, *Pvalb* levels did vary in both DG and neocortex as a function of SD (values plotted as cumulative distributions in **Figure 7**). These changes moved in opposite directions, with DG neurons showing SD-driven decreases in *Pvalb* labeling intensity (**Figure 7A**), and neocortex showing SD-driven increases in *Pvalb* (**Figure 7D**). However, mean *Pvalb* intensity values (calculated per area) were not affected by SD in either IEG+ PV+ interneurons or IEG-PV+ interneurons, in any structure (**Figure S3**).

**Figure 7.**
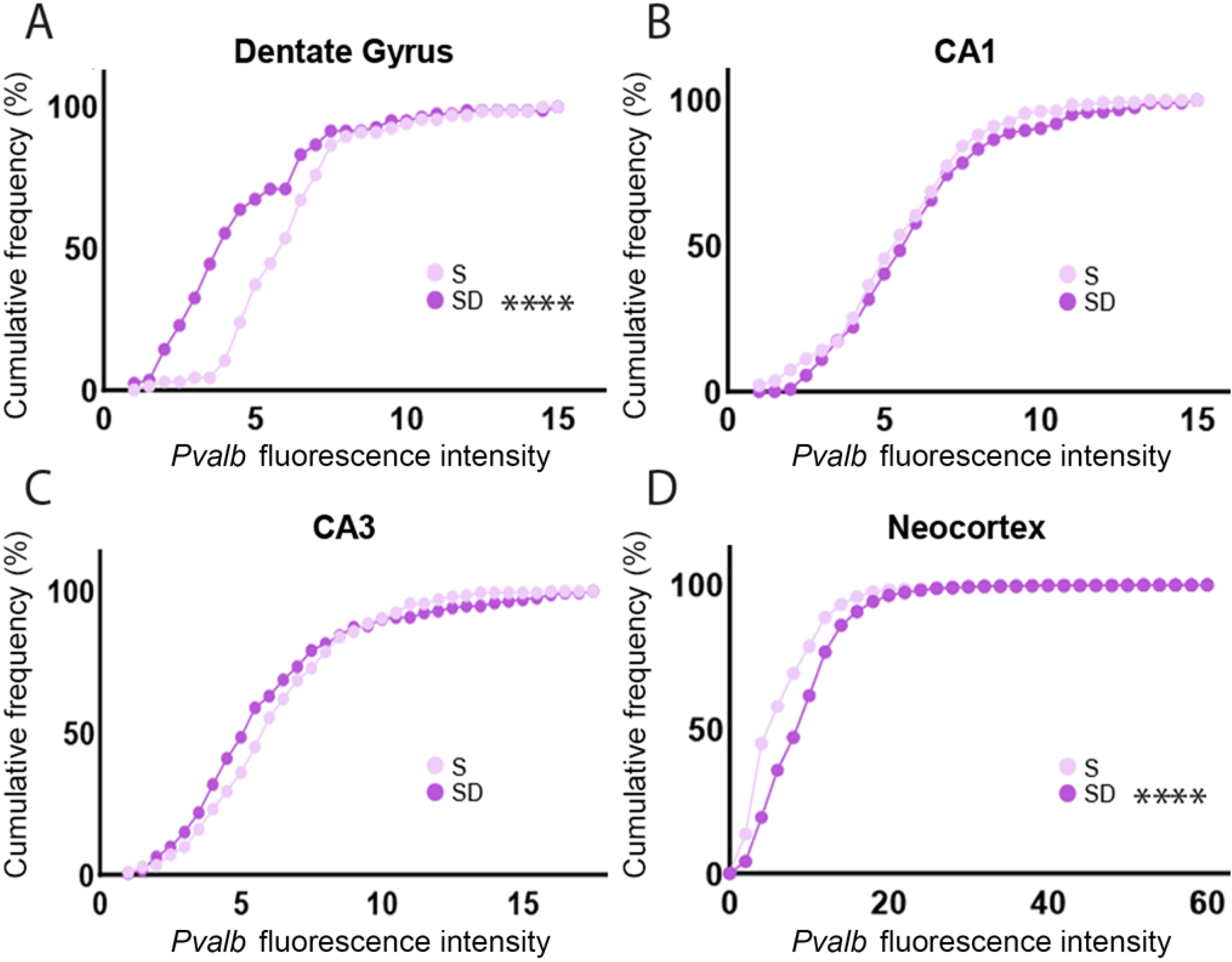
SD alters neuronal *Pvalb* mean fluorescence intensity in a region- and subregion-specific manner. Cumulative frequency distributions showing the impact of 6-h SD on *Pvalb* fluorescence intensity in *Pvalb+* cells of the hippocampus and neocortex. **(A)** 6-h SD significantly decreased mean fluorescence intensity of *Pvalb* within *Pvalb+* cells of the DG while having no significant effect on **(B)** CA1 or **(C)** CA3 intensity. **(D)** 6-h SD significantly increased mean fluorescence intensity of *Pvalb* within *Pvalb+* cells of the neocortex. Hippocampal (DG, CA1, CA3) and neocortical bin widths for cumulative frequency distributions set at 0.5 and 2 respectively; **** indicates *p* < 0.0001, Holm–Sidak *post hoc* test vs. Sleep.

Together these data suggest that SD drives relatively modest changes in *Homer1a*, *Arc*, and *Cfos* in neocortical PV+ interneurons, but does not affect these transcripts in hippocampal PV+ interneurons, and that SD drives differential hippocampal vs. neocortical changes in expression of *Pvalb*.

### Cell type- and region-specific effects of SD on ribosome-associated transcripts involved in circadian timekeeping

SD has previously been implicated in regulating core molecular clock genes’ expression. As is true for IEG expression, the extent to which SD differentially impacts core clock gene expression as a function of cell type and regions is unclear. Consequently, we quantified ribosome-associated transcript abundance for core clock genes-*Clock, Per1, Per2, Cry1, Cry1, and Bmal1-* after SD in Camk2a+ neurons and PV+ interneurons of the neocortex and hippocampus (**Figure 8**). Consistent with findings from whole neocortical tissue (Franken et al., 2007; Hoekstra et al., 2019), we found that 3-h SD significantly increased *Per2* expression in neocortical Camk2a+ neurons and PV+ interneurons (**Figure 8A**). In contrast, SD had no significant impact on transcript abundance in the hippocampus of either population. Longer-duration (6-h) SD resulted in no further changes in neocortical transcript abundance (with *Per2* levels tending to remain elevated in both Camk2a+ neurons and PV+ interneurons) (**Figure 8B**). Within the hippocampus, 6-h SD significantly altered abundance of ribosome-associated *Per2, Cry1*, and *Cry2* transcripts in Camk2a+ neurons (increasing *Per2* and *Cry1*, decreasing *Cry2*), while having no significant effect on transcript abundance in PV+ interneurons.

**Figure 8.**
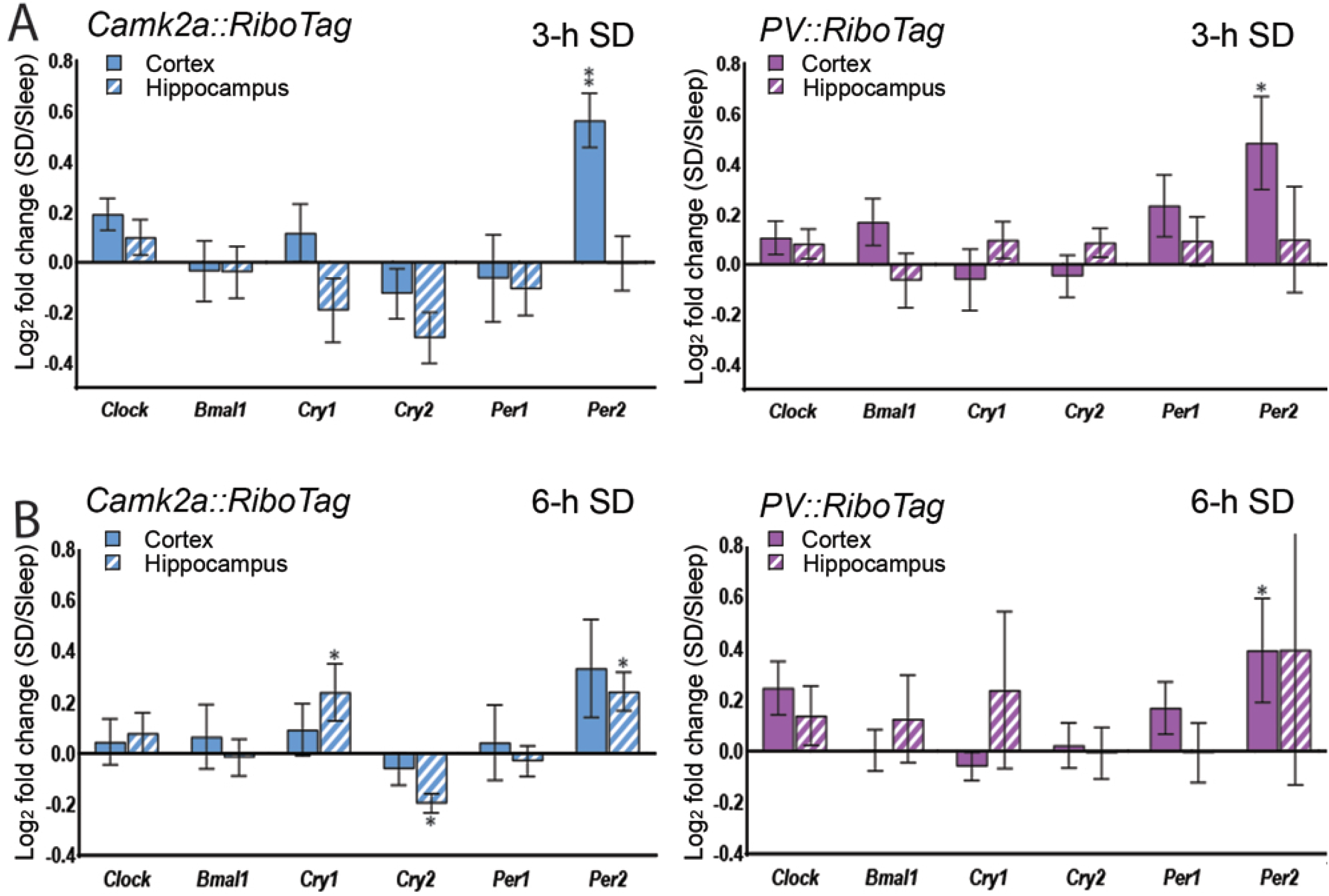
SD alters ribosome-associated transcripts encoding core clock genes in a cell type and region-specific manner. **(A)** 3-h SD significantly increased *Per2* abundance on ribosomes in Camk2a+ (blue) and PV+ (magenta) neocortical neurons; no significant changes in core clock transcripts were observed in hippocampal neurons. **(B)** After 6-h SD, *Per2* abundance remained significantly elevated in neocortical PV+ interneurons. Ribosome-associated *Cry1*, *Cry2*, and *Per2* were all altered after 6-h SD in the hippocampal Camk2a+ neuron population. No significant change observed among PV+ interneurons. Transcript level changes are presented as a log_2_ fold change between SD and *ad lib* sleep mice. Values indicate mean ± SEM with propagated error; * and ** indicate *p* < 0.05 and *p* < 0.01, respectively, Holm–Sidak *post hoc* test vs. Sleep.

We also quantified (after SD vs Sleep) the abundance of ribosome-associated mRNAs encoding other cellular timekeeping components: *Rev-Erbα, Dbp, Ted, Nfil3, and Dec1* (**Figure 9**). We found significant heterogeneity in how these auxiliary clock genes responded to SD in different cell types and regions. None of the transcripts were significantly altered in either cell type in the hippocampus, with either 3-h or 6-h SD (**Figure 9A-B**). However, within the neocortex, both 3-h and 6-h SD significantly increased cortical *Nfil3* and *Dec1* abundance in PV+ interneurons. While these transcripts were not significantly altered in neocortical Camk2a+ neurons, 6-h SD significantly decreased *Rev-Erbα* expression in Camk2a+ neocortical neurons (**Figure 9B**).

**Figure 9.**
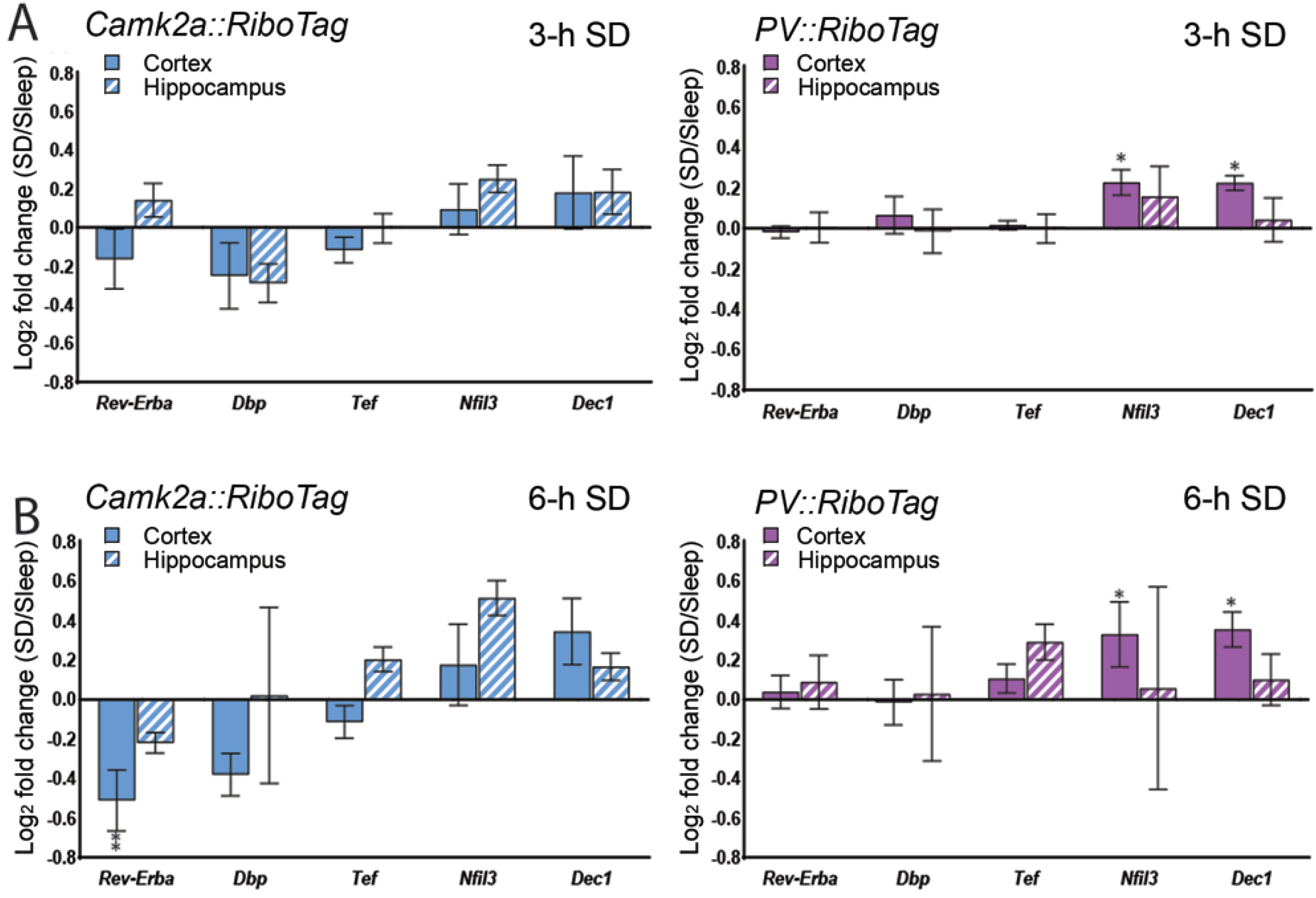
SD differentially alters circadian clock modifiers in Camk2a+ and PV+ neocortical populations. **(A)** 3-h SD had no significant effect on ribosome-associated circadian clock modifier transcripts among Camk2a+ (blue) neurons in neocortex, but increased *Nfil3* and *Dec1* expression among neocortical PV+ interneurons (magenta). **(B)** 6-h SD significantly decreased *Rev-Erbα* abundance on ribosomes in Camk2a+ neocortical neurons. No transcripts were significantly altered by SD in either neuron population in hippocampus. Transcript level changes are presented as a log_2_ fold change between SD and *ad lib* sleep mice. Values indicate mean ± SEM with propagated error; * and ** indicate *p* < 0.05 and *p* < 0.01, respectively, Holm–Sidak *post hoc* test vs. Sleep.

## Discussion

Here, using TRAP, we have identified SD-driven molecular changes unique to specific cell populations in hippocampus and neocortex. Numerous studies have used transcriptome (Cirelli et al., 2004; Vecsey et al., 2012) or proteome (Noya et al., 2019; Poirrier et al., 2008) profiling of these structures following sleep vs. SD as a way of clarifying the functions of sleep in the brain. We find that comparing across structures, there are large differences in SD effects on ribosome-associated transcripts. For example, while even brief (3-h) SD increases abundance of plasticity-mediating transcripts in neocortical Camk2a+ neurons and PV+ interneurons (**Figure 2**) few of these transcripts are altered in hippocampus even after longer SD. This is particularly true for hippocampal PV+ interneurons, for which none of the transcripts are significantly altered by SD. SD-driven changes in abundance for IEG transcription regulators follow a similar pattern (**Figure 3**), with hippocampal PV+ interneurons in particular being refractory to SD. Our *in situ* analysis of mRNA abundance in PV+ and PV-neurons (**Figures 4–6**) is consistent with this interpretation, and suggests that even within neocortex, SD-driven changes in these transcripts’ abundance are relatively modest in PV+ interneurons (**Figure 4**).

While IEGs are generally assumed to reflect specific patterns of recent neuronal activity (Tyssowski and Gray, 2019), there are brain region- and microcircuit-specific differences in IEG expression which reflect neurons’ network connectivity patterns (Gonzalez et al., 2019; Tyssowski et al., 2018). Moreover, IEG expression in PV+ interneurons is regulated by distinct cellular pathways and is differentially gated by neuronal activation (Cohen et al., 2016). Indeed, some studies have failed to detect IEGs in PV+ interneurons altogether (Imamura et al., 2011; Vazdarjanova et al., 2006), and our present results showing relatively low expression in the PV+ interneuron population (**Figures 4–6**). However, insofar as abundance of all of these transcripts is regulated by neuronal activity to some degree (Donato et al., 2013; Yap and Greenberg, 2018), our present data support two broad conclusions. First, neuronal activation in the hippocampus is reduced relative to neocortex during SD. Second, PV+ interneuron activity may vary less as a function of SD than Camk2a+ neuron activity.

The former conclusion has major implications for the field of learning and memory, where pronounced and selective effects of sleep disruption on hippocampal processes (e.g., episodic and spatial memory consolidation) have been well described (Havekes and Abel, 2017; Puentes-Mestril et al., 2019; Saletin and Walker, 2012). In hippocampal structures such as the DG and CA1, available data suggest that both markers of neuronal activity and synaptic plasticity are disrupted after SD (Delorme et al., 2019; Havekes et al., 2016; Ognjanovski et al., 2018; Raven et al., 2019; Tudor et al., 2016). Our present data largely confirm these findings, and suggest that particularly in DG and CA1 (**Figure 5**), there is little evidence of neuronal activation during SD. Indeed, we find that DG neurons show decreased *Pvalb* expression after SD, while neocortical neurons simultaneously show increased expression (**Figure 7**). Critically, *Pvalb* expression levels have been shown to correlate with both PV+ interneuron activity level and the relative amounts of excitatory to inhibitory input PV+ interneurons receive (Donato et al., 2013). Thus we conclude that SD increases excitatory input to PV+ interneurons in neocortex, while simultaneously decreasing excitatory input to DG. This conclusion parallels our recent work showing differential effects of SD on another activity marker, *Arc*, in DG vs. neocortex, and suggests that SD may have a uniquely disruptive effect on network activity in DG.

The latter conclusion also has important implications for maintenance of excitatory-inhibitory (E-I) balance during SD. Recent data suggest that E-I balance normally varies over the course of the day, in a sleep-dependent manner (Bridi et al., 2020). Furthermore, prior evidence from both whole-tissue transcriptome profiling and immunohistochemistry has suggested that SD may differentially affect connections from excitatory to inhibitory neurons (and vice versa) in structures like the neocortex (Del Cid-Pellitero et al., 2017; Puentes-Mestril and Aton, 2017). Because sleep loss is one of the major risk factors for triggering seizure onset in epilepsy (Frucht et al., 2000; Lawn et al., 2014), an underlying mechanism might be differential activation of, or plasticity in, interneurons vs. principal neurons with SD. Interactions between PV+ interneurons and principal neurons are particularly important in both regulation of attention (Aton, 2013) and in generating network oscillations important for memory consolidation (Ognjanovski et al., 2018; Ognjanovski et al., 2017). Insofar as SD may disrupt both attention and memory consolidation, differential effects on activity of PV+ and Camk2a+ neurons in the hippocampus and neocortex may be an important underlying mechanism.

Because many of the transcripts quantified here (e.g., *Arc*, *Homer1a*, *Narp*, and *Bdnf*) play a critical role in activity-regulated synaptic plasticity, the fact that their abundance on translating ribosomes in Camk2+ and PV+ neurons is differentially altered by SD (**Figure 2**) also has intriguing implications. For example, it suggests that SD could lead to long-lasting changes in the E-I balance and information processing capacity of neocortical and hippocampal circuits. This may be a plausible mechanism for some of the reported longer-lasting brain metabolic (Wu et al., 2006) and cognitive (Belenky et al., 2003; Chai et al., 2020; Dinges et al., 1997) effects of SD (i.e., those that do not normalize with recovery sleep).

Alterations in brain clock gene expression with SD has been widely reported (Franken et al., 2007; Mongrain et al., 2011; Wisor et al., 2002; Wisor et al., 2008). Along with transcripts such as *Homer1a* (Maret et al., 2008; Zhu et al., 2020), SD-driven increases in transcripts such as *Per2* are hypothesized to play a role in homeostatic aspects of sleep regulation (Franken et al., 2007; Mang and Franken, 2015). Our data suggest that similar to plasticity-regulating transcripts (including *Homer1a*), SD-mediated changes in clock gene transcripts on ribosomes are cell type- and brain region-specific (**Figures 8** and **9**). For example, while *Per2* increases on both Camk2a+ and PV+ neocortical neuron-derived ribosomes with as little as 3 h SD, no clock gene transcripts are altered in the hippocampus with 3-h SD (**Figure 8A**, **Figure 9A**). Another example is *Rev-erbα*, which is significantly reduced after 6-h SD, but only in neocortical Camk2a+ neurons. An interesting and important issue, raised by our findings, is that SD-driven changes in particular core clock transcripts’ abundance do not move in the same direction, as they normally would during a 24-h cycle (e.g., *Cry1*, *Cry2*, and *Per2*; **Figure 8**). This suggests that SD-driven changes in these transcripts may not be driven by canonical E-box elements, consistent with recent findings (Mongrain et al., 2011). However, because changes in these transcripts may have numerous downstream effects on transcription of other clock-control genes (Chiou et al., 2016; Schmutz et al., 2010), these SD-driven changes may have even more numerous downstream effects that changes in plasticity effectors’ transcripts. Future studies will be needed to quantify longer-term cell type-specific changes to physiology and structure initiated during SD, and the molecular events responsible for these changes.

Together our data suggest that effects of SD on plasticity, timekeeping, and homeostatic regulation of brain circuitry is heterogeneous, and likely involves subtle modifications to microcircuits (e.g., those in hippocampal subregions and neocortical layers) critical for appropriate brain function.

## Acknowledgements

The authors are grateful to members of the Aton lab, and to Drs. Natalie Tronson, Geoff Murphy, and Michael Sutton for helpful feedback on this manuscript. This work was supported by research grants from the NIH (DP2 MH 104119) and the Human Frontiers Science Program (N023241-00_RG105) to SJA.

## Supplementary material

### Supplementary table

**Table S1.**
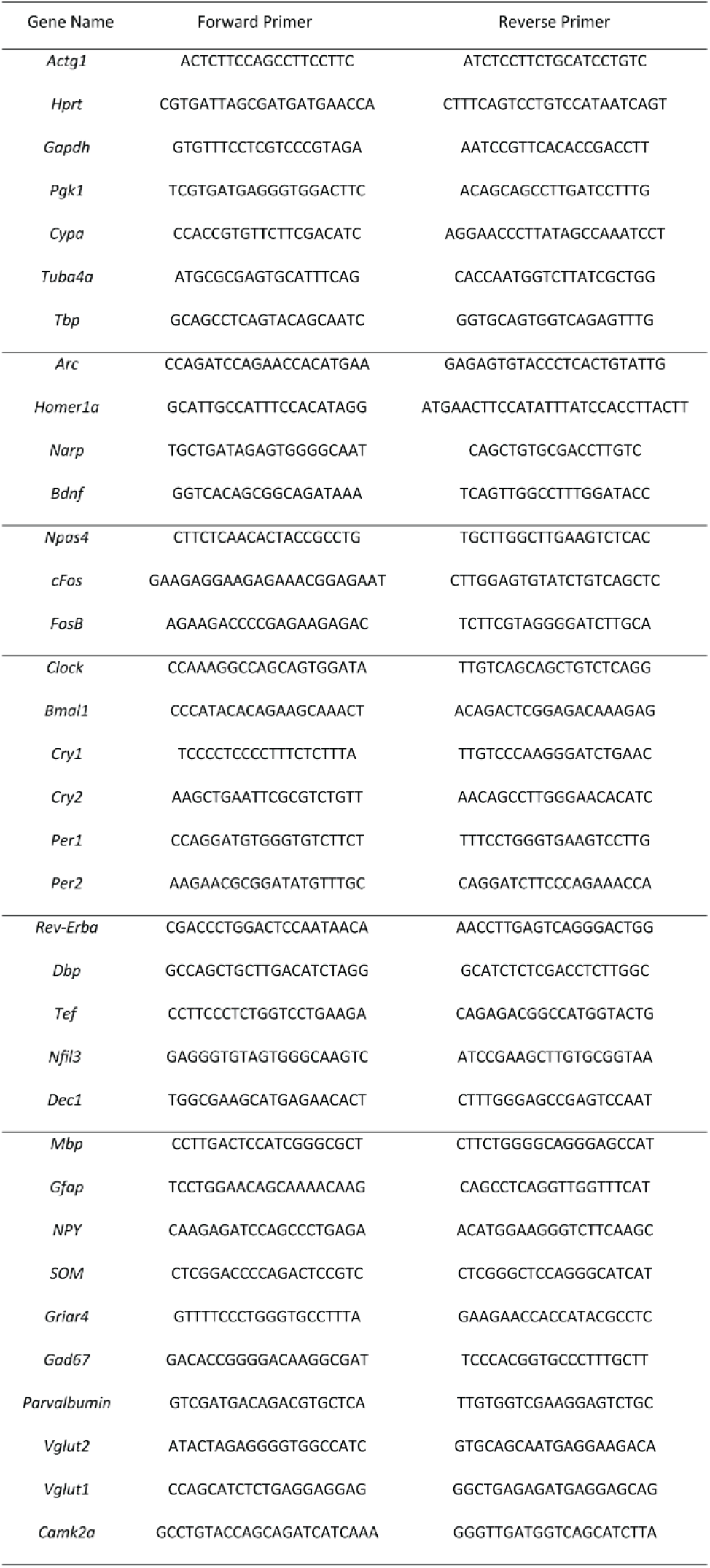
Primer Designs.

**Table S2.**
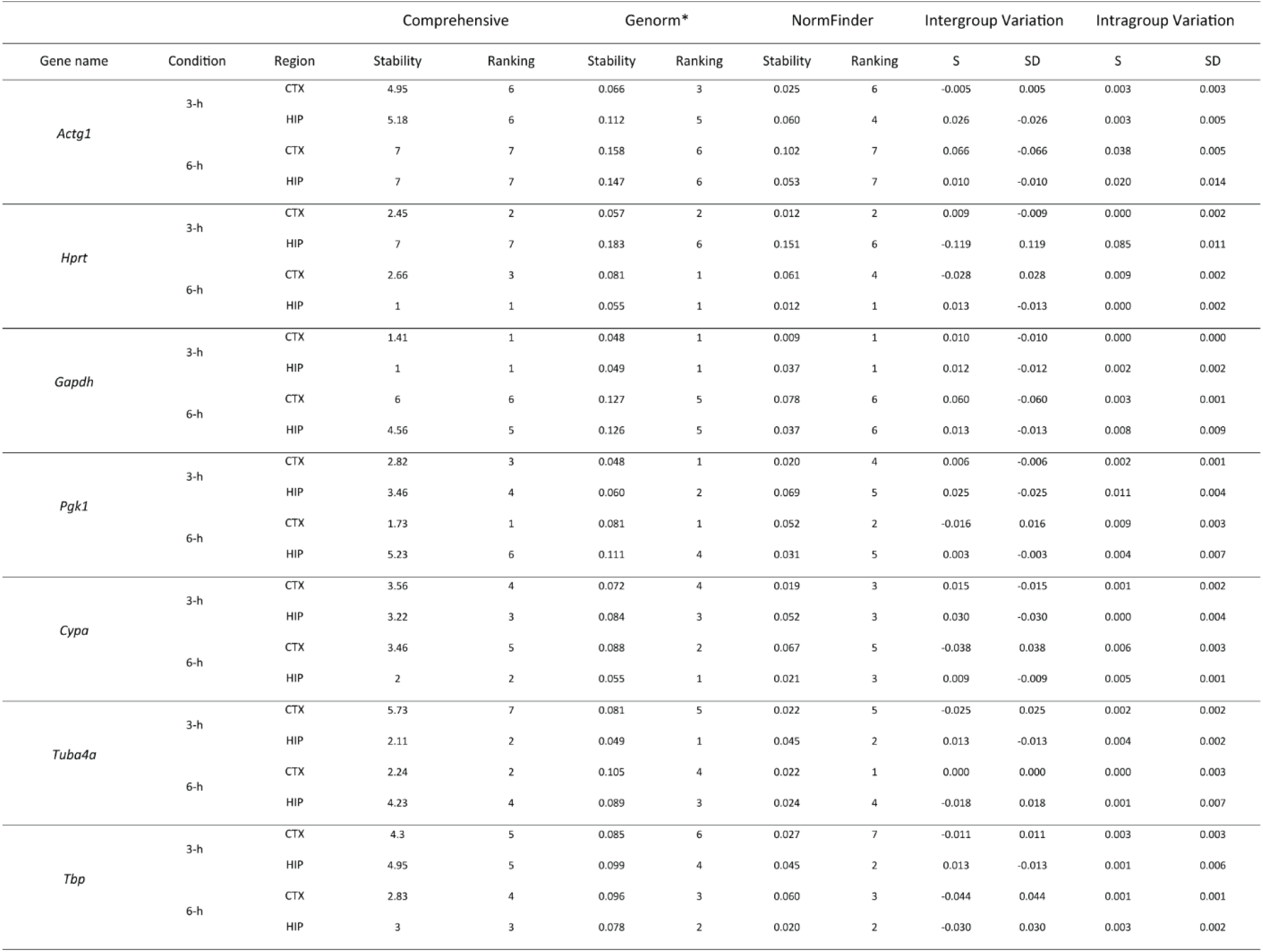
Housekeeping stability analysis for Camk2a:RiboTagsamples. *Genorm automatically calculates the stability measure for the two most stable genes.

**Table S3.** Housekeeping stability analysis for PV:RiboTag samples. *Genorm automatically calculates the stability measure for the two most stable genes.

| Gene name | Condition | Region | Comprehensive |  | Genorm* |  | NormFinder |  | Intergroup Variation |  | Intragroup Variation |  |
| --- | --- | --- | --- | --- | --- | --- | --- | --- | --- | --- | --- | --- |
|  |  |  | Stability | Ranking | Stability | Ranking | Stability | Ranking | S | SD | S | SD |
| *Actg1* | 3-h | CTX | 5.12 | 5 | 0.089 | 6 | 0.033 | 5 | -0.007 | 0.007 | 0.003 | 0.003 |
|  |  | HIP | 3.76 | 5 | 0.189 | 5 | 0.046 | 4 | -0.032 | 0.032 | 0.028 | 0.001 |
|  | 6-h | CTX | 7 | 7 | 0.142 | 6 | 0.063 | 7 | -0.032 | 0.032 | 0.019 | 0.010 |
|  |  | HIP | 3.56 | 4 | 0.144 | 5 | 0.033 | 3 | 0.025 | -0.025 | 0.003 | 0.012 |
| *Hprt* | 3-h | CTX | 5.44 | 6 | 0.080 | 4 | 0.035 | 7 | 0.030 | -0.030 | 0.000 | 0.001 |
|  |  | HIP | 6.74 | 7 | 0.246 | 7 | 0.100 | 7 | -0.046 | 0.046 | 0.031 | 0.059 |
|  | 6-h | CTX | 1.19 | 1 | 0.077 | 1 | 0.033 | 3 | 0.020 | -0.020 | 0.002 | 0.001 |
|  |  | HIP | 1.57 | 1 | 0.106 | 1 | 0.030 | 2 | 0.022 | -0.022 | 0.005 | 0.005 |
| *Gapdh* | 3-h | CTX | 1.68 | 1 | 0.041 | 1 | 0.032 | 4 | -0.020 | 0.020 | 0.001 | 0.002 |
|  |  | HIP | 3.46 | 4 | 0.166 | 4 | 0.040 | 3 | -0.015 | 0.015 | 0.010 | 0.005 |
|  | 6-h | CTX | 2.51 | 3 | 0.099 | 3 | 0.032 | 2 | 0.023 | -0.023 | 0.001 | 0.001 |
|  |  | HIP | 2.34 | 3 | 0.106 | 1 | 0.019 | 1 | 0.039 | -0.039 | 0.000 | 0.008 |
| *Pgk1* | 3-h | CTX | 2.28 | 3 | 0.041 | 1 | 0.031 | 3 | -0.019 | 0.019 | 0.001 | 0.003 |
|  |  | HIP | 1.32 | 1 | 0.143 | 1 | 0.014 | 1 | -0.015 | 0.015 | 0.002 | 0.000 |
|  | 6-h | CTX | 2.28 | 2 | 0.077 | 1 | 0.031 | 1 | -0.001 | 0.001 | 0.008 | 0.002 |
|  |  | HIP | 4.16 | 5 | 0.135 | 3 | 0.042 | 4 | -0.012 | 0.012 | 0.016 | 0.007 |
| *Cypa* | 3-h | CTX | 6 | 7 | 0.085 | 5 | 0.034 | 6 | 0.021 | -0.021 | 0.002 | 0.003 |
|  |  | HIP | 6.24 | 6 | 0.215 | 5 | 0.068 | 6 | 0.027 | -0.027 | 0.056 | 0.004 |
|  | 6-h | CTX | 4.56 | 5 | 0.12 | 5 | 0.045 | 5 | -0.025 | 0.025 | 0.003 | 0.009 |
|  |  | HIP | 7 | 7 | 0.242 | 6 | 0.085 | 6 | -0.070 | 0.070 | 0.002 | 0.138 |
| *Tuba4a* | 3-h | CTX | 1.86 | 2 | 0.059 | 2 | 0.016 | 1 | -0.003 | 0.003 | 0.000 | 0.000 |
|  |  | HIP | 2.78 | 3 | 0.152 | 2 | 0.061 | 5 | 0.045 | -0.045 | 0.010 | 0.024 |
|  | 6-h | CTX | 5.23 | 6 | 0.112 | 4 | 0.047 | 6 | 0.036 | -0.036 | 0.004 | 0.006 |
|  |  | HIP | 6 | 6 | 0.177 | 5 | 0.052 | 5 | -0.008 | 0.008 | 0.043 | 0.002 |
| *Tbp* | 3-h | CTX | 3.56 | 4 | 0.07 | 3 | 0.025 | 2 | -0.003 | 0.003 | 0.005 | 0.000 |
|  |  | HIP | 1.86 | 2 | 0.143 | 1 | 0.019 | 1 | 0.004 | -0.004 | 0.012 | 0.000 |
|  | 6-h | CTX | 3.72 | 4 | 0.091 | 2 | 0.039 | 4 | -0.020 | 0.020 | 0.002 | 0.005 |
|  |  | HIP | 1.86 | 2 | 0.124 | 2 | 0.019 | 1 | 0.004 | -0.004 | 0.003 | 0.001 |

### Supplementary figure legends

**Figure S1.**
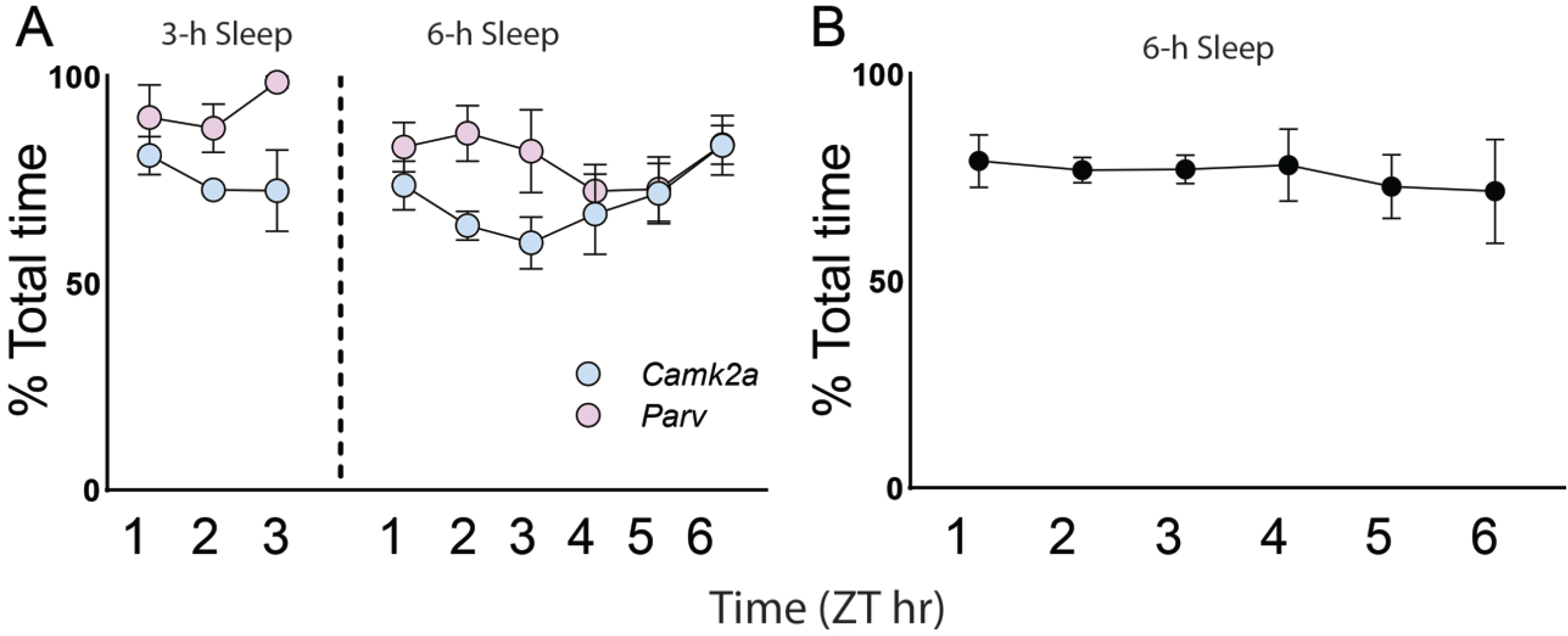
Total sleep time in freely-sleeping mice. Proportion of time spent in *ad lib* sleep between ZT0 and ZT3 or ZT6 for mice used in RT-qPCR experiments **(A)** and *in situ hybridization* experiments **(B)**. Sleep behavior (during which mice were observed to be inactive and in stereotyped sleep posture) was quantified in 5-min or 2-min intervals across the *ad lib* sleep period (for 6-h and 3-h experiments, respectively). Values expressed as a mean percentage of total time spent in sleep (± SEM), in 60-min intervals.

**Figure S2.**
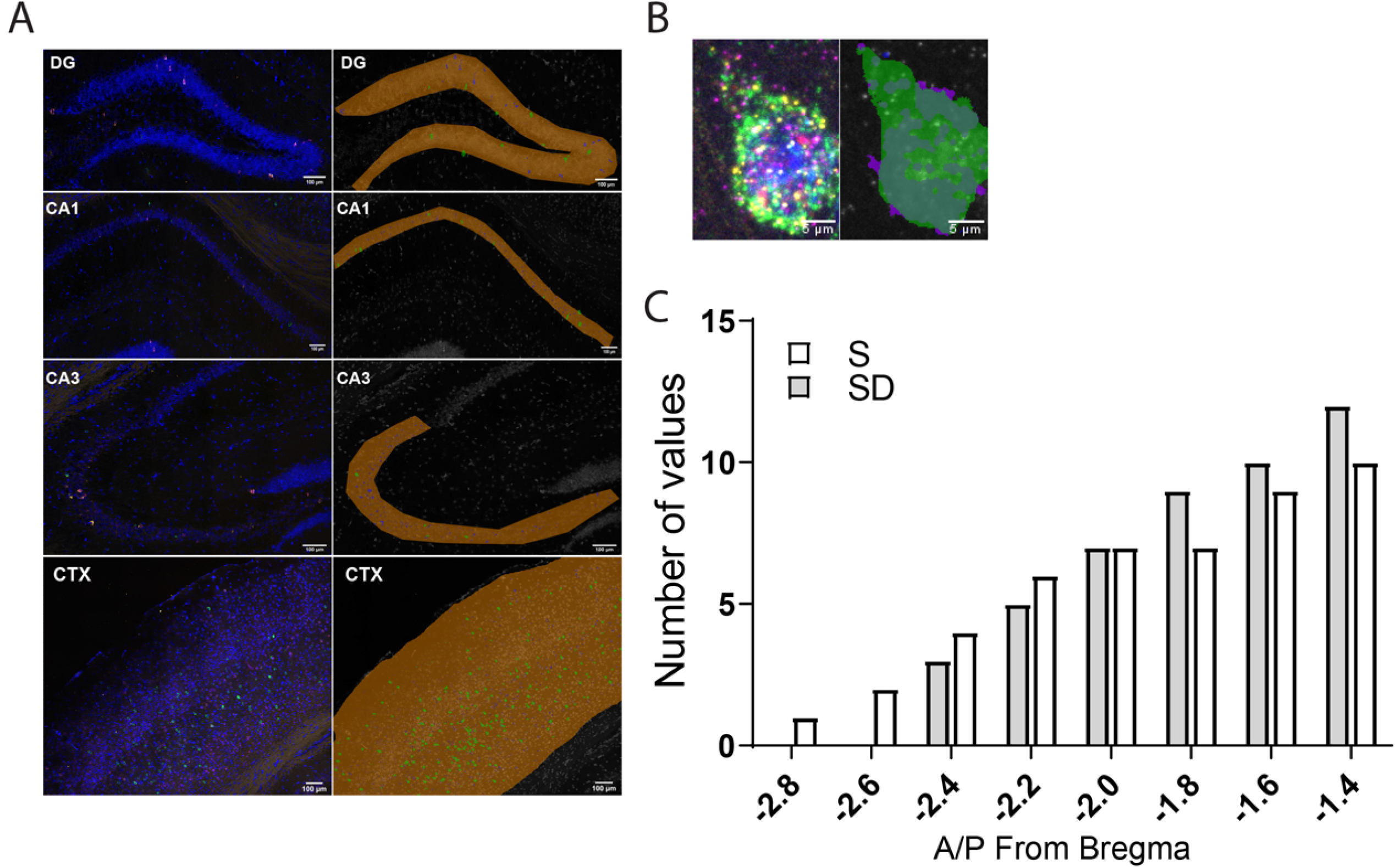
Strategy for quantification of fluorescence *in situ* signals. **(A)** Anatomical regions for quantification were demarcated manually (shown in orange). Within these anatomical regions, *Pvalb* (green) fluorescence delineated PV+ and non-PV+ ROIs. Background was defined as any area not expressing IEG (red) fluorescence. An automated protocol then calculated the total fluorescence intensity and area of each ROI and background area. These values were used to estimate the number of IEG fluorescence dots within each ROI. **(B)** Example of IEG and *Pvalb* fluorescence and identification. **(C)** Cumulative frequency distribution of A/P coordinates (relative to Bregma) for brain sections used in analysis.

**Figure S3.**
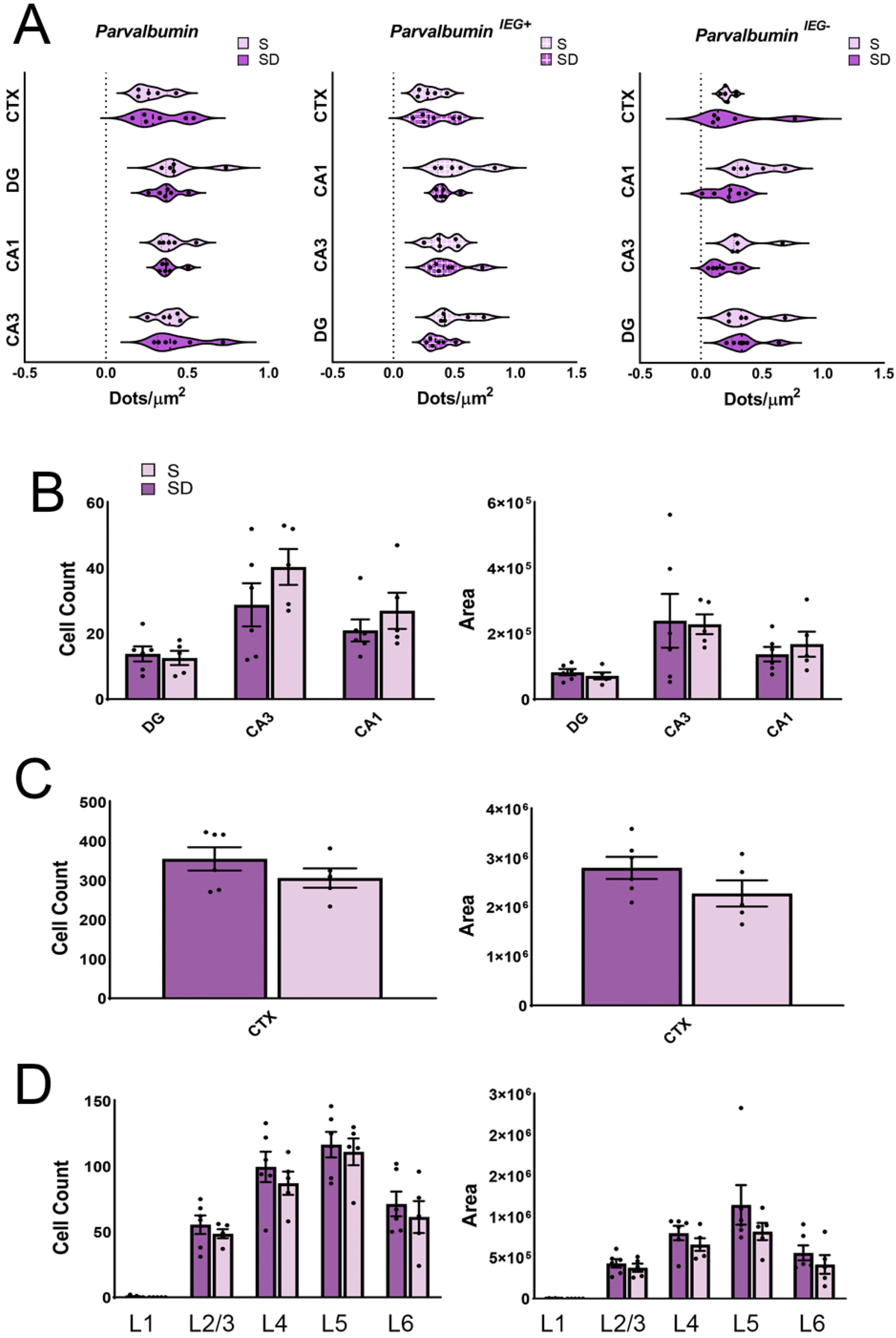
Mean *Pvalb* mRNA expression is similar between freely-sleeping and SD mice. **(A)** Mean *Pvalb* expression levels were similar in sleeping and SD mice, in all, IEG+, and IEG-PV+ interneurons. **(B-D)** Neither the total *Pvalb*+ cell count nor the *Pvalb*+ area differed between sleeping and SD mice, for either **(B)** hippocampal areas DG, CA1, or CA3, **(C)** whole neocortex, **(D)** or cortical layers 1-6. Values indicate mean ± SEM; *N.S.*, Holm–Sidak *post hoc* test vs. Sleep.

